# DNA segment capture by Smc5/6 holo-complexes

**DOI:** 10.1101/2022.10.09.511515

**Authors:** Michael Taschner, Stephan Gruber

## Abstract

Three distinct SMC complexes facilitate chromosome folding and segregation in eukaryotes, presumably by DNA translocation and loop extrusion. How SMCs interact with DNA is however not well understood. Among the SMC complexes, Smc5/6 has dedicated roles in DNA repair and in preventing a lethal buildup of aberrant DNA junctions. Here, we describe the reconstitution of ATP-dependent topological DNA loading by Smc5/6 rings. By inserting cysteine residues at selected protein interfaces, we obtained covalently closed compartments upon chemical cross-linking. We show that two SMC subcompartments and the kleisin compartment topologically entrap a plasmid molecule, but not the full SMC compartment. This is explained by a looped DNA segment inserting into the SMC compartment with the kleisin neck gate locking the loop in place when passing between the two DNA flanks and closing. This DNA segment capture strictly requires the Nse5/6 loader, which opens the neck gate prior to DNA passage. Similar segment capture events without gate opening may provide the power stroke for DNA translocation/loop extrusion in subsequent ATP hydrolysis cycles. Our biochemical experiments thus offer a unifying principle for SMC ATPase function in loading and translocation/extrusion, which is likely relevant to other members of the family of SMC proteins too.

## Introduction

In eukaryotes, three distinct Structural Maintenance of Chromosomes (SMC) complexes (cohesin, condensin, and Smc5/6) share essential tasks in the maintenance of chromosome structure and the faithful transmission of genetic information during nuclear division [reviewed in ^1^]. Some of these ATP-powered DNA-folding machines are known to form large DNA loops by loop extrusion ^2–4^. This is thought to allow condensin to compact chromatids in mitosis and cohesin to participate in gene regulation, DNA repair and recombination in interphase. However, loop extrusion or merely DNA translocation may well be a conserved feature of all pro- and eukaryotic relatives. Smc5/6 has indeed recently been shown to extrude DNA loops or translocate along DNA *in vitro* (Pradhan et al., 2022), but the cellular functions and relevance of these activities remain unclear. DNA entrapment has long been considered a basic and essential feature of SMC function ^1^. It provides a simple explanation for how SMC complexes stay *in cis* (on a given DNA molecule) over time, maintain directionality of translocation, and readily bypass obstacles up to few tens of nm in size that they encounter frequently on the chromosomal translocation track. However, the requirement of DNA entrapment and the exact nature of the DNA interaction remain contested with all possibilities being considered in the recent literature (**Fig 1A**): topological DNA entrapment, DNA loop entrapment (also denoted as pseudo-topological entrapment), and exclusively external DNA association. The latter scenario was bolstered by the perceived bypass of very large obstacles by purified cohesin and condensin in single molecule imaging experiments ^5^. Cryo-electron microscopy studies have not yet been able to adequately address this question and particularly little is known about how Smc5/6 associates with DNA. Resolving DNA topology will be vital to refute and refine models for how the SMC ATPase powers DNA translocation and loop extrusion ^6,7^.

**Figure 1:**
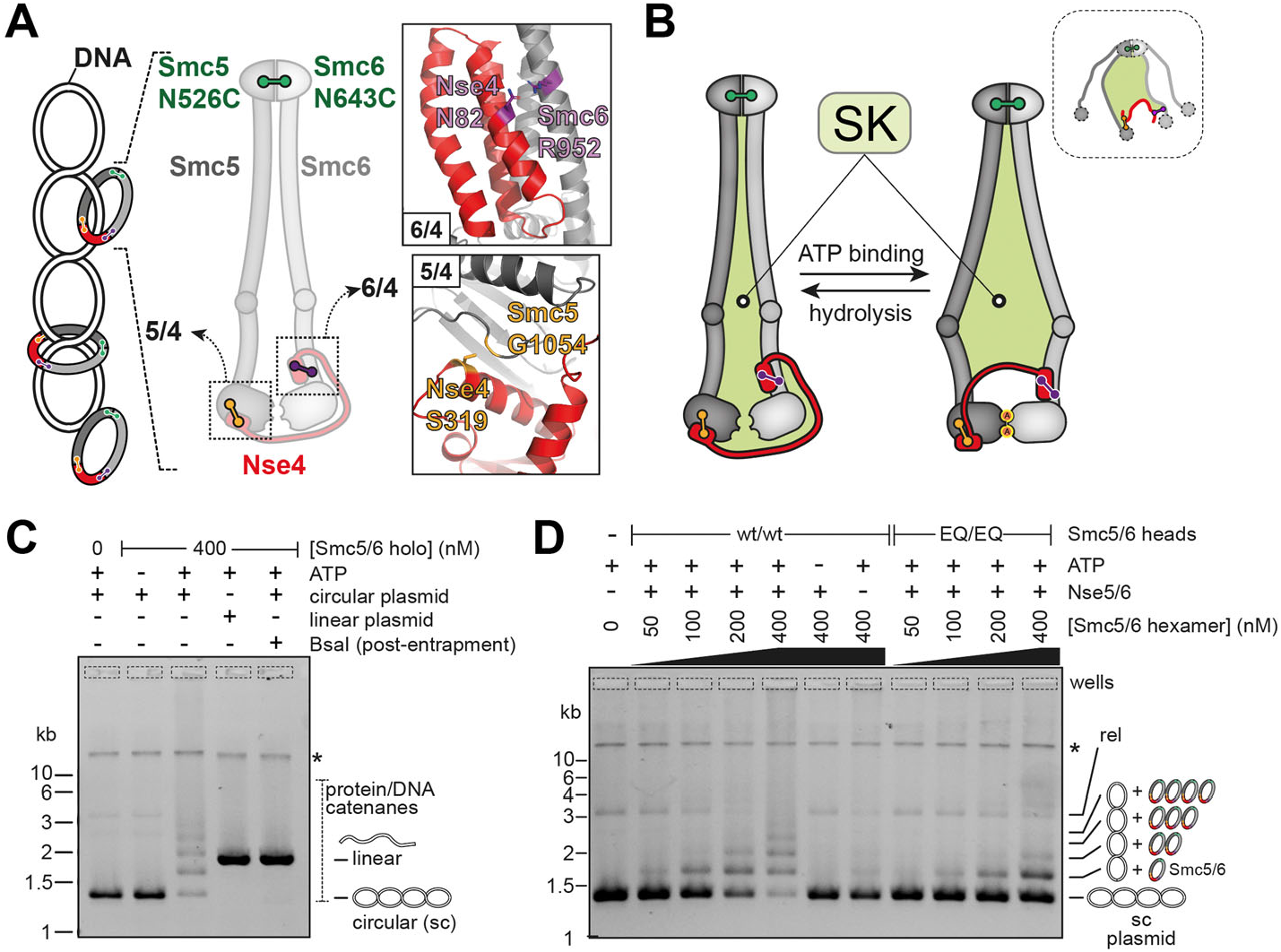
DNA entrapment in Smc5/6. **(A)** Schematic view of a supercoiled circular DNA substrate (left) bound to SMC/kleisin rings (with topological DNA entrapment, top; non-topological DNA entrapment, middle; no DNA entrapment, bottom) and of the Smc5/6 ring subunits (middle panel). Positions of cysteines engineered for cross-linking of the hinge, Smc6/Nse4 and Smc5/Nse4 interfaces are indicated in green, purple, and orange colors, respectively. Close-up view of the positions of cysteines at the Smc6/Nse4 (top right panel) and Smc5/Nse4 (bottom right panel) interfaces. **(B)** Two schematic representations of the SMC/Kleisin (SK) compartment depending on the positioning of the Nse4 kleisin subunit. Three cysteine pairs—necessary to covalently close it—are indicated with colored handlebars. A cartoon showing the cross-linked complex with the colored compartment after denaturation is shown in the dashed box. Positions of hinge as well as N- and C-terminal ATPase domains are still shown as half-ovals and circles, respectively. **(C)** Co-isolation of cross-linked Smc5/6 proteins with plasmid DNA by agarose gel electrophoresis. Results obtained with octameric Smc5/6 holo-complex harboring cysteine pairs for cross-linking of the SK ring, comparing linear and circular DNA substrates. Traces of contaminating linear DNA from *E. coli* are marked with an asterisk. **(D)** As in (C) but with titration of the protein complex and comparison of entrapment efficiencies between the wildtype and ATP-hydrolysis-deficient (EQ/EQ) complexes. Controls without either ATP or the Nse5/6 loader are also included. The plasmid substrate is largely supercoiled (sc), but a relaxed (rel) form is also visible.

Complete loss of Smc5/6 function leads to severe chromosome segregation defects in mitosis and meiosis ^8–12^. Toxic DNA structures such as unresolved recombination intermediates, DNA intertwinings, or incompletely replicated chromosomal regions prevent proper chromosome segregation in the absence of Smc5/6 ^13–15^. DNA translocation by Smc5/6 may be required to detect and eliminate such DNA junctions. The Smc5/6 holo-complex comprises eight subunits: Smc5-6 and Nse1-6 in *S. cerevisiae* (**fig S1A**). All of them cause lethality when removed by gene disruption ^9^. Smc5-6 and Nse4 together form the conserved ATPase core with an elongated shape that is characteristic for all SMC complexes ^16–21^. Smc5 and Smc6 proteins harbor the canonical ‘hinge’ domains for heterotypic dimerization connected via ~35 nm long antiparallel coiled-coil ‘arms’ to globular ABC-type ‘head’ domains with highly conserved motifs for ATP-binding and hydrolysis ^22–24^. The kleisin subunit Nse4 bridges the head of the κ-SMC protein Smc5 to the head-proximal coiled coil (‘neck’) of the v-SMC protein Smc6. This creates the SMC/kleisin (SK) ring structure shown in other SMC complexes to be in principle capable of entrapping DNA ^25–29^. Nse4 serves as attachment point for two kite proteins (the Nse1 and Nse3 subunits), also found in the prokaryotic SMC complexes ^30^, while the Smc5/6-specific subunit Nse2 attaches to the coiled-coil arm of Smc5 ^31^. Finally, a stable sub-complex is formed by the Nse5 and Nse6 subunits ^18,20^. It contacts the Smc5/6 hexamer via multiple interfaces including one on the arms and one on the heads. Single-molecule tracking and biochemical studies suggest a function of Nse5/6 in chromosome loading ^20,32^. In *S. pombe* (but not in *S. cerevisiae*), disruption of *nse5* and *nse6* genes cause less severe phenotypes when compared to deletion of other Smc5/6 subunits, indicating that the hexamer can partially function without Nse5/6 in fission yeast ^33^. Curiously, loop extrusion by *S. cerevisiae* Smc5/6 does not depend on the loader Nse5/6 and is actually inhibited by it, implying that Smc5/6 has essential functions beyond loop extrusion ^34^. A dedicated loader appears to be involved in viral restriction by Smc5/6 in human cells ^35^.

The ATP hydrolysis cycle involves major structural rearrangements in the SMC complex ^36–41^. Three main conformations have been delineated for Smc5/6 (**fig S1B**) ^16–21^: (1) In the ‘ATP-engaged state’, two molecules of ATP are sandwiched by residues of the Walker A and B motifs of the Smc5 head and the signature motif of the Smc6 head and *vice versa*. The engaged heads keep the two head-proximal arms at a distance, thus opening an SMC compartment ^36,42,43^. (2) In the ‘juxtaposed state’, the heads are ATP-disengaged and the Smc5 and Smc6 arms co-align yielding a closed SMC compartment ^21^. (3) The ‘inhibited state’ relies on the intercalation of the loader Nse5/6 between the arms and heads of Smc5/6 ^19,20^, thus preventing head engagement and ATP hydrolysis. The inhibition is in turn overcome by the binding of a suitable DNA substrate ^20^. A head/DNA interface on top of the engaged heads was described for several SMC complexes ^38,44-50^. Such a DNA-clamping state has recently also been described for Smc5/6 by cryo-electron microscopy ^43^.

Here we demonstrate that Smc5/6 holo-complexes efficiently entrap DNA molecules. Using the reconstituted DNA loading reaction, we elucidate the position and topology of DNA in the Smc5/6 complex, and we unambiguously identify the neck gate as DNA entry gate.

## Results

### Topological DNA entrapment by the Smc5/6 ring

Salt-stable DNA binding by the octameric yeast Smc5/6 holo-complex depends on a circular DNA substrate, being indicative of a topological component of the Smc5/6 DNA association ^20^. Here we delineate the position of DNA in the Smc5/6 complex by creating covalently closed DNA compartments followed by protein-DNA co-isolation under protein-denaturing conditions (**Fig 1A**). First, we tested whether the perimeter of the Smc5/6 ring entraps plasmid DNA by cross-linking the three ring interfaces to generate a covalently closed SK ring. We used structure prediction (by AlphaFold-Multimer) to position cysteine pairs at two ring interfaces (**Fig 1A**) ^51,52^. We observed robust chemical cross-linking of the Smc5/Nse4 interface (~80 %) and the Smc6/Nse4 interface (~90 %) upon addition of BMOE (**fig S2A**, lanes 3 and 4), as shown for the Smc5/6 hinge ^20^.

When cysteine pairs at all three ring interfaces were combined, cross-linking gave rise to a significant fraction of covalently closed Smc5/6 SK ring species (**fig S2A**, lane 8) (**Fig 1B**). The cysteine mutations did not have strong effects on the Smc5/6 ATPase activity (**fig S2B**). We then separated protein-DNA catenanes from free DNA species by agarose gel electrophoresis (schematics in **fig S3A**) ^53^. Cross-linking at the three SK ring interfaces resulted in a characteristic laddering of a small (1.8 kb) plasmid at an elevated Smc5/6 concentration (400 nM) (**Fig 1C**), presumably due to the reduced mobility of DNA molecules associated with increasing numbers of Smc5/6 complexes. The laddering was fully dependent on ATP and disappeared when plasmid DNA was linearized by a restriction enzyme prior to gel electrophoresis, as expected for a topological interaction (**Fig 1C**). This loading reaction reached a steady state relatively quickly (within minutes; see below). Moreover, three protein preparations lacking one of the six engineered cysteines showed little or no DNA laddering, with the residual gel shift likely explained by low levels of off-target cross-linking (**fig S2C**).

The laddering was virtually abolished in the absence of the loader Nse5/6 (**Fig 1D**) and also clearly reduced by ATP-hydrolysis deficient mutations in both Smc5 and Smc6 (‘EQ/EQ’), indicating that ATP hydrolysis promotes more robust DNA entrapment. Based on similar experiments using protein gel electrophoresis, we estimate that a large fraction of Smc5/6 entraps plasmid DNA under our experimental conditions (**fig S2D**). We conclude that topological DNA loading by the Smc5/6 holo-complex is a robust and efficient reaction.

### DNA entrapment in the kleisin compartment

We next wondered where DNA might be located in the Smc5/6 complex. We previously showed that under the conditions promoting salt-stable DNA binding and DNA entrapment (**Fig 1C-D**) the complex exhibits efficient head engagement as measured by cysteine cross-linking ^20^. Head engagement produces an SMC (‘S’) and a kleisin (‘K’) compartment (see schemes in **Fig 2A**). DNA must be in at least one of the two compartments. We first combined the cysteine pair at ATP-engaged heads with cysteines at both SMC/Nse4 interfaces to generate a covalently closed K compartment, which yielded robust DNA entrapment again in an ATP and loader-dependent manner (**Fig 2B**, top panel). The efficiency of entrapment in the K compartment appeared somewhat reduced when compared to the SK ring but this is likely explained by the reduced efficiency of engaged-heads cross-linking [~25 %; ^20^]. A series of samples cross-linked at different time points after mixing showed that entrapment was detectable after 30 seconds and was saturated within a few minutes in both the SK ring and the K compartment (**fig S2E**), supporting the notion that the two types of entrapment emerge from the same biochemical reaction and possibly correspond to the same state. We conclude that the K compartment of the ATP-engaged Smc5/6 holocomplex is occupied by DNA.

**Figure 2:**
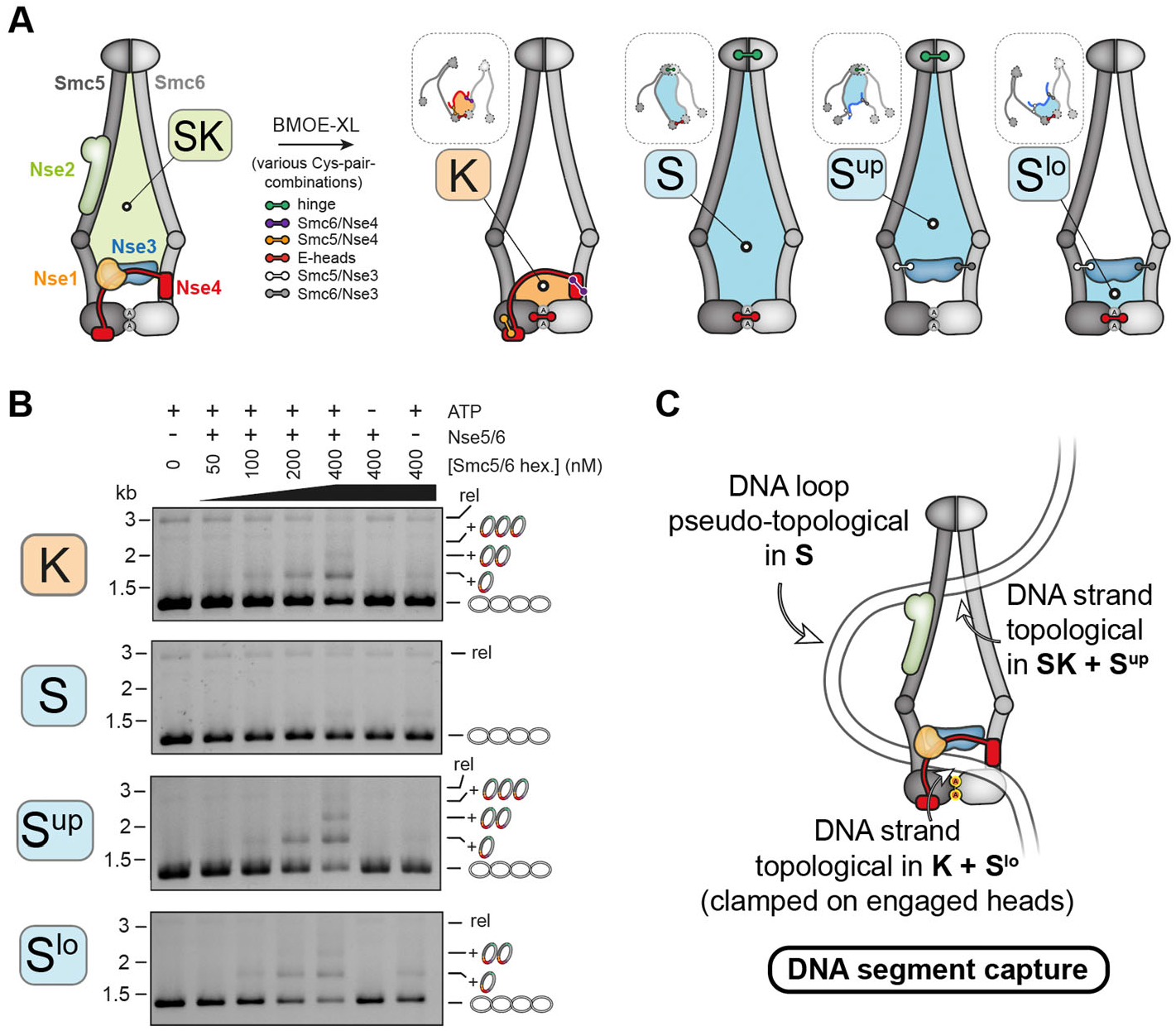
DNA entrapment in Smc5/6 sub-compartments. **(A)** Schematic representation of sub-compartments formed during ATP-dependent head engagement. The scheme on the left shows the complete Smc5/6 hexamer with the SK compartment highlighted as in Fig 1. Combinations of Cys-pairs (colored handlebars as indicated) lead to covalent closure of kleisin (K) and SMC (S) compartments, with the latter being split into upper (S^up^) and lower (S^lo^) compartments by Nse3 crosslinking as indicated. Note that the schemes on the right only show the cross-linked subunits that remain attached after denaturation. Schemes in dashed boxes indicate compartments after denaturation as in (**Fig 1B**). **(B)** Co-isolation of cross-linked Smc5/6 proteins with plasmid DNA by agarose gel electrophoresis. Results obtained with protein preparations harboring cysteine pairs for cross-linking of the K (top panel), S (second panel), S^up^ (third panel), or S^lo^ compartments (bottom panel). As in **Fig 1D. (C)** Schematic representation of a DNA segment capture state explaining the findings in (B).

### DNA segment capture in the SMC compartment

Next, we combined head and hinge cysteines to create a covalently closed S compartment (**Fig 2A**). Contrary to the K compartment, no loader-dependent DNA entrapment was observed for the S compartment (**Fig 2B**, second panel). This may suggest that the S compartment is devoid of DNA when heads are ATP-engaged. Alternatively, a DNA loop (rather than a single DNA double helix) may thread into the S compartment with the pseudo-topologically held DNA being lost upon protein denaturation even after cross-linking. To avoid confusion with the tobe-extruded DNA loop, we designate this configuration as DNA segment capture (**Fig 2C**, right scheme) ^54^. To test for DNA segment capture, we sought to split the S compartment into two sub-compartments that each entrap a single DNA double helix rather than a DNA loop. A cryoEM structure showed that the Nse3 subunit may split the S compartment into halves by contacting the Smc5 as well as the Smc6 coiled coil in a DNA clamping state ^43^. We generated cysteine pairs to cross-link Nse3 to Smc5 and to Smc6 (**fig S4**) and combined them with hinge cysteines to create an upper SMC compartment (S^up^) and with the head cysteines to generate a lower SMC compartment (S^lo^) (**Fig 2A**). Intriguingly, the cysteine combinations for the S^up^ and the S^lo^ compartments both resulted in robust DNA laddering (**Fig 2B**, lower panels), demonstrating that a DNA segment is efficiently captured in the SMC compartment and held as a loop with one DNA passage in the S^lo^ compartment and another one in the S^up^ compartment (**Fig 2C**).

### The loader Nse5/6 opens the neck gate in Smc5/6

In the course of the above experiments, we noticed that the cross-linking of the cysteine pair at the Smc6/Nse4 interface, designated as neck gate, is strongly sensitive to ligands. While cross-linking was robust (~ 90%) with or without ATP and DNA in the absence of the loader (**Fig 3**, lanes 2-5), it was strongly hampered when the loader Nse5/6 was added, especially when plasmid DNA was present (**Fig 3**, lanes 7 and 8). Of note, such a behavior was not observed for cysteine pairs at the hinge ^20^, along the coiled-coil arms (**fig S5**) or at the Smc5/Nse4 interface (**fig S6A**). However, addition of the loader led to some off-target crosslinking of a native cysteine in Nse5 with Smc5(G1054C) (**fig S6B**). More importantly, however, neck gate cross-linking in the octamer was partially restored by addition of ATP alone (**Fig 3**, lane 9) and virtually fully restored by addition of both ATP and DNA (lane 10). Similar cross-linking efficiencies were observed with complexes harboring all cysteines for SK ring cross-linking (**fig S6C**). Experiments performed with complexes carrying mutations of active site residues showed that this stimulatory effect of DNA on neck gate closure requires ATP-engagement of Smc5/6 heads (**fig S6D**, lanes 9 and 10) but not ATP hydrolysis (**fig S6E**, lanes 9 and 10). Altogether, the presented results thus suggest that binding of the Smc5/6 hexamer to the loader Nse5/6 specifically detaches one of the three ring interfaces. DNA clamping, *i*.*e*., ATP-engagement of SMC heads in conjunction with DNA binding at the head/DNA interface, appears to promote gate closure. Interestingly, fusion of Nse4 to Smc6, but not to Smc5, is lethal in *S. cerevisiae*, indicating that opening of the neck gate is essential for Smc5/6 function (**fig S6F** and ^17^). The neck gate is largely closed in the presence of ATP and DNA, implying that once loaded onto DNA, Smc5/6 holo-complexes form a ring that encompasses the DNA double helix.

**Figure 3:**
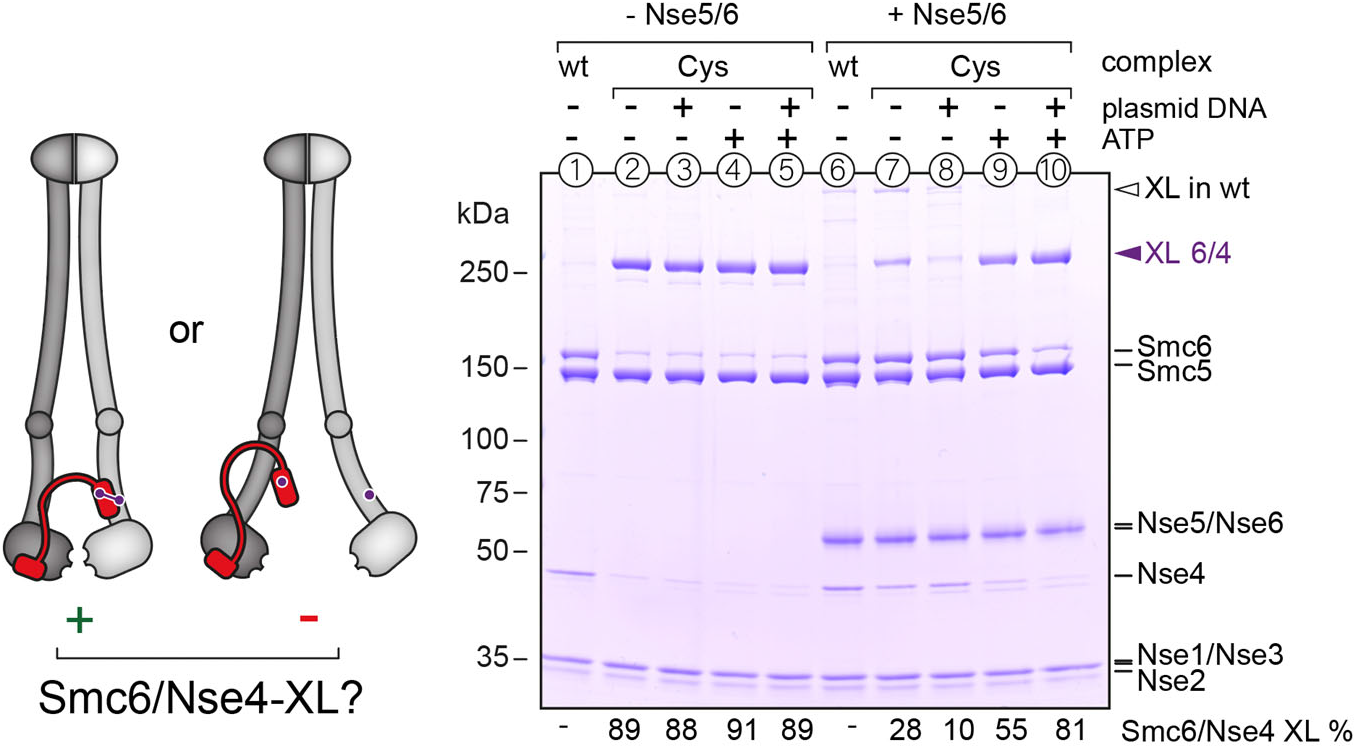
Opening of the neck gate in Smc5/6. Cross-linking of purified Smc5/6 hexamers with cysteines at the Smc6/Nse4 interface in the presence and absence of ligands. Detection of cross-linked species by SDS-Page and Coomassie staining. Loss of cross-linking suggests that the gate is open. Equivalent experiments for this interface containing ATPase head mutations as well as for other cysteine pairs are shown in **fig S6**.

### DNA clamping is a prerequisite for efficient gate closure and DNA entrapment

The above observations suggested that DNA binding plays a role in neck gate closure. We next determined whether the head/DNA interface is indeed crucial for topological DNA entrapment or important only subsequently for example for the conversion of a putative ‘DNA holding’ intermediate into the DNA segment capture state (see later). Based on sequence conservation and structure comparison (**fig S7 A-C**), we identified three positively charged residues on Smc5 (K89, R139, R143) and another three on Smc6 (R135, K200, K201) as putative DNA binding residues (**Fig 4A**). To test whether these residues are important for DNA entrapment, we mutated them by alanine and glutamate substitution (charge removal and reversal, respectively) in isolation or in combination. Mutant alleles were tested for functional complementation of a respective deletion mutant in yeast using plasmid shuffling ^55^. *smc5* alleles harboring double or triple glutamate substitutions resulted in a strong growth phenotype, while alanine mutations were apparently tolerated well (**Fig 4B**). The *smc6* gene was somewhat more sensitive to the mutagenesis with the triple alanine mutant also exhibiting clear growth retardation (**Fig 4C**). A selection of residues based on a recent cryo-EM map ^43^ resulted in similar outcomes (**fig S7D** and **E**). The deficiencies of the single, double, and triple alanine mutants in *smc6* were aggravated when combined with the *smc5(3A)* allele, suggesting that these residues have partially overlapping functions (**fig S7F**) as previously observed for bacterial Smc ^38^.The sextuple alanine mutant strain [*smc5(3A), smc6(3A)* or 3A3A] displayed a null-like phenotype, supporting the notion that the head/DNA interface is crucial for an essential function of Smc5/6.

**Figure 4:**
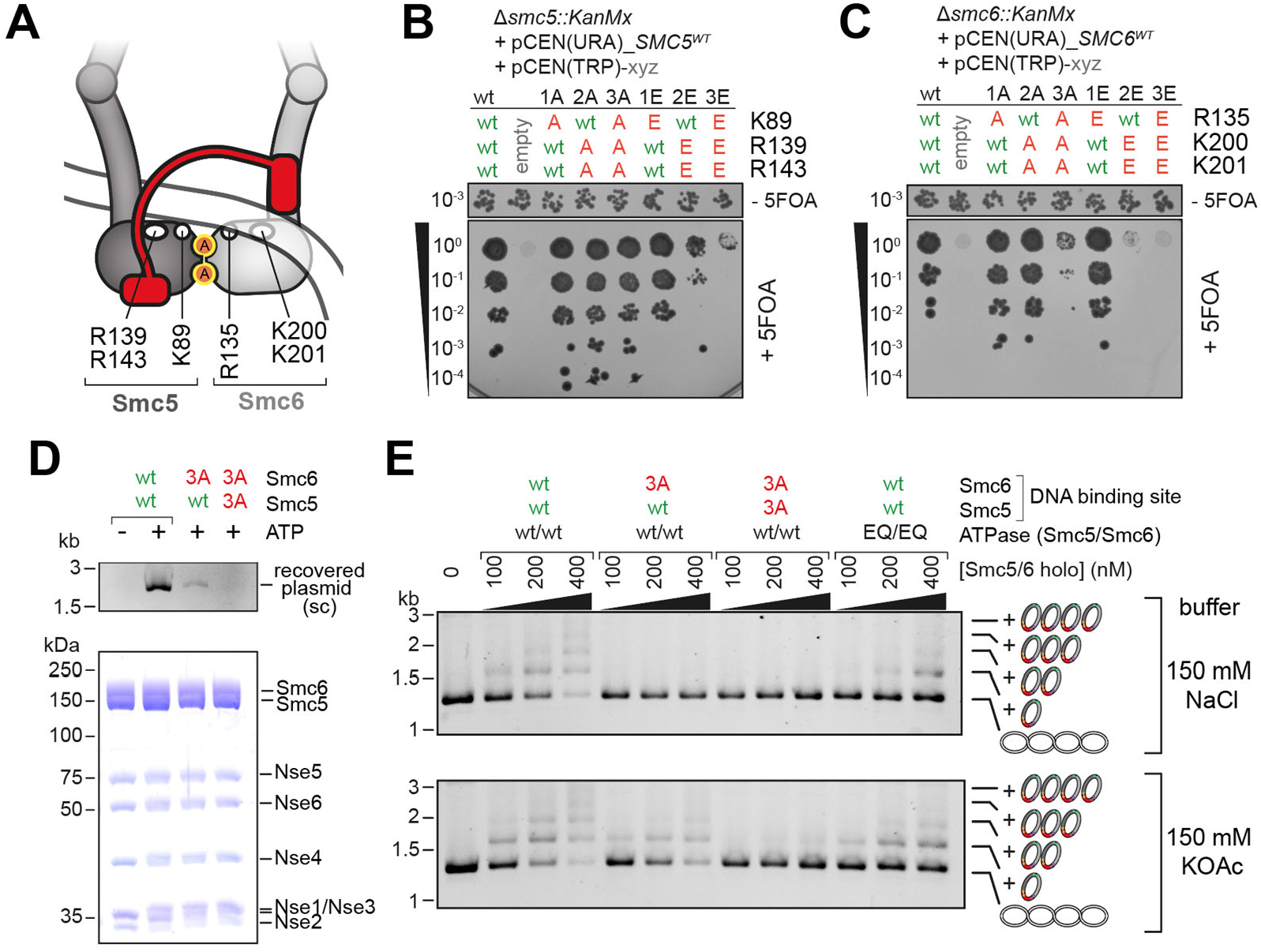
DNA clamping is essential for DNA entrapment. **(A)** The positions of the selected residues are schematically displayed for the DNA clamping state. See also **fig S7A. (B)** Positively charged residues on the Smc5 head were mutated to alanine (‘A’) or glutamate (‘E’) in isolation or in combination as indicated. The mutant alleles were tested for function by plasmid shuffling. Counterselection against a pCEN(URA3) plasmid carrying a wildtype *SMC5* allele by addition of 5-fluorouracil (‘5FOA’) revealed *smc5* mutants resulting in a growth defect. **(C)** As in (B) for residues in Smc6. **(D)** Salt-stable DNA binding with wild-type and mutant Smc5/6 as determined by protein immobilization using a Twin-Strep tag on Smc6 (bottom panel) and detection of co-isolated plasmid DNA (‘recovered plasmid’) by agarose gel electrophoresis (top panel). **(E)** DNA entrapment in the SK ring of wild-type, DNA-binding, and ATP-hydrolysis defective Smc5/6 mutants. Using standard DNA entrapment salt conditions (150 mM NaCl) as in **Fig 1D** (top panel) or buffer with reduced ionic strength (150 mM KOAc) as used for the salt-stable DNA binding assay in (D).

Selecting one partially and one fully defective mutant (*smc6(3A)* and 3A3A, respectively) we tested for DNA loading of Smc5/6 when DNA binding at the heads is compromised. First, we measured ATP-dependent salt-stable DNA binding by pull-down assays as described previously [^20^; **fig S3B**]. The Smc6(3A) variant only poorly recovered plasmid DNA, while the 3A3A variant was completely unable to do so, together strongly suggesting that DNA binding is crucial for Nse5/6 mediated DNA loading (**Fig 4D**). Next, we combined the Smc6(3A) and the 3A3A mutations with the cysteine variants for SK cross-linking and DNA entrapment. Using the standard buffer conditions (150 mM NaCl), both mutants failed to support any DNA entrapment (**Fig 4E**, top panel). When the reaction buffer was adjusted to mimic the conditions used for salt-stable DNA binding (150 mM KOAc; a ‘milder’ salt with larger ions), the Smc6(3A) variant supported residual DNA entrapment, while the 3A3A variant did not (**Fig 4E**, bottom panel). These results demonstrate that the head/DNA binding interface is crucial for topological DNA entrapment by Smc5/6. Of note, the mutant complexes showed slightly reduced ATPase activity (**fig S2B**). However, the defect in DNA entrapment is not explained by the lack of ATP hydrolysis, since the EQ/EQ complex quite efficiently entrapped DNA despite failing to support any noticeable ATPase activity (**Fig 4E and fig S2B**). Moreover, the mutant complexes produced ring crosslinking albeit at somewhat reduced efficiency, in particular the 3A3A mutant, an effect that was more pronounced in the stringent salt buffer (**fig S6G**). The latter is likely explained by defects in neck gate closure, as the complex lacking DNA binding residues on Smc5 and Smc6 heads also failed to display DNA-stimulated neck gate closure (**fig S6H**). DNA clamping might thus serve (at least) two related purposes during DNA entrapment: guiding kleisin over clamped DNA prior to closure and promoting gate closure itself.

### DNA passage through the neck gate

We wondered whether DNA might indeed enter the Smc5/6 ring by passing through the neck gate that opens upon association with the loader Nse5/6 and closes upon DNA encounter. To test this hypothesis directly, we created an Smc6-Nse4 fusion protein. The linker peptide covalently connects the C-terminus of the Smc6 polypeptide to the N-terminus of Nse4 and includes a recognition sequence for cleavage by the 3C protease (**fig S8A**). This construct is non-functional *in vivo* (**fig S6F;** as previously reported for a related construct {Serrano, 2020 #98}). The reconstituted Smc5/6 complex harboring this fusion protein hydrolyzed ATP normally (**fig S2B**). We next combined the Smc6-Nse4 fusion protein with cysteines for DNA entrapment experiments (see schematics in **fig S8B**). When combining cysteines at the hinge and the Smc5/Nse4 interface with the Smc6-Nse4 fusion protein, the ring species failed to produce any gel shift, demonstrating that DNA entrapment is not possible when Smc6 is linked to Nse4, as expected if the neck gate indeed serves as DNA entry gate (**Fig 5A**, lanes 5-7 and **fig S8B**). Curiously, however, we recovered DNA laddering when combining the Smc6-Nse4 fusion with only the neck gate cross-link albeit with an altered laddering pattern with distinctively smaller steps, probably owing to the smaller size of the protein ring species formed with Smc6 and Nse4 protein only (**Fig 5A**, lanes 8-10 and **fig S8B**). Combining the Smc6-Nse4 fusion with all three cysteine pairs yielded DNA laddering with larger as well as smaller laddering steps presumably due to partial cross-linking (**Fig 5A**, lanes 11-13 and **fig S8B**). To reconcile these observations, we suggest that the linker may be long enough to embrace the incoming DNA, thus not blocking the loading reaction itself (**Fig 5B**). This would lead to pseudo-topological entrapment of a DNA loop in an expanded ‘SK-plus-linker’ compartment (see schematics of closed and denatured compartments in **Fig 5A**), thus explaining the absence of laddering when the Smc6-Nse4 interface was only closed by the peptide linker but not by the cross-link. Cleavage of the linker peptide by 3C protease before (**fig S8C**) or after (**Fig 5C**) the loading reaction eliminated the entrapment by the linker compartment and also removed the smaller-sized steps observed with the fusion protein plus all three cysteine pairs. Also consistent with the linker wrapping around DNA, we find that the Smc6-Nse4 fusion complex supported normal or near-normal salt-stable DNA binding regardless of whether the peptide linker was left intact or cleaved, whereas a control complex with an open kleisin did not (**fig S8D**). Taken together, these results strongly suggest that the engineered peptide linker indeed embraces the incoming DNA and identify the neck gate as DNA entry gate.

**Figure 5:**
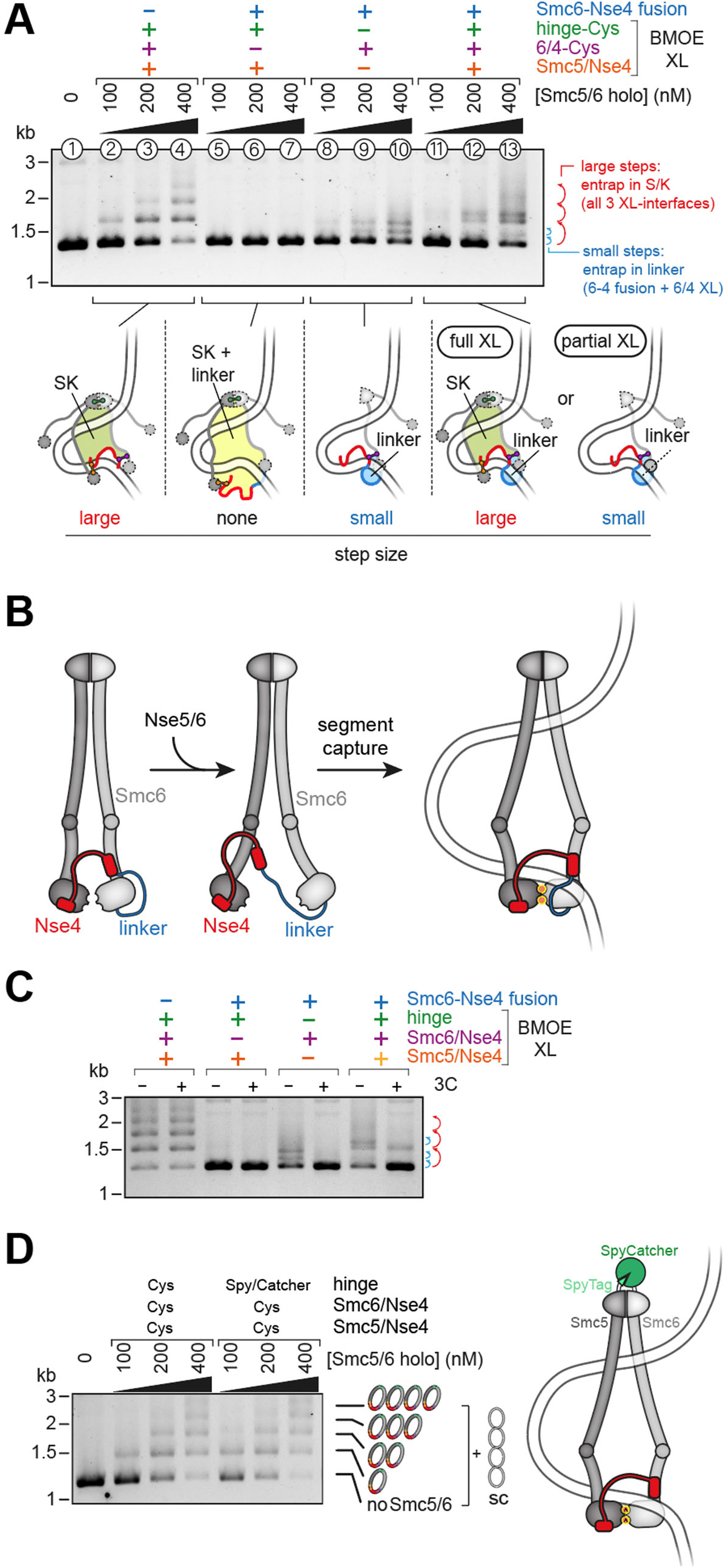
DNA passes through the Smc6/Nse4 gate. Co-isolation of cross-linked Smc5/6 proteins with plasmid DNA by agarose gel electrophoresis. Results obtained with protein preparations containing combinations of Cys-pairs and an Smc6-Nse4 fusion protein. Schematic drawings of crosslinked and denatured protein species are shown below the gel, with the closed compartments indicated. Different observed stepsizes of DNA ladders are caused by differences in the size of the crosslinked protein species. **(B)** Schematic representation of DNA clamping in the presence of an Smc6-Nse4 fusion protein. The length of the linker allows it to wrap around the DNA strand clamped on ATP-engaged heads. **(C)** As in (A) but with only a high concentration of complex, and with or without post-entrapment opening of the linker using the HRV 3C protease. **(D** Co-isolation of cross-linked Smc5/6 proteins with plasmid DNA by agarose gel electrophoresis, comparing a protein preparation harboring cysteine pairs for cross-linking of the SK ring with one in which the hinge Cysteine-pair is replaced by a Spytag/Spy-Catcher fusion. Topological DNA entrapment in Smc5/6 is not prevented by the hinge-fusion, unlike shown for cohesin.

Our experiments imply that all loading happens via the neck gate. However, a recent study showed that (in addition to the neck gate) the hinge serves as DNA entry gate for topological entrapment by cohesin *in vitro* ^53^. Related experiments performed with covalently linked Smc5/6 hinge domains (using a Spy-tag/Spy-Catcher pair, **fig S8E**) did not display any DNA entrapment defects (**Fig 5D**) confirming that passage via the hinge does not notably contribute to DNA entrapment in Smc5/6 and highlighting intriguing differences in DNA loading between these SMC complexes.

### The Smc5/6 ring entraps plasmid DNA *in vivo*

Finally, we wanted to investigate whether DNA entrapment by the Smc5/6 complex also occurs *in vivo*. We introduced the cysteine pairs for SK ring crosslinking into the *nse4, smc5* and *smc6* genes in budding yeast by allelic replacement. Following BMOE cross-linking in cells, immunoprecipitation and Southern blotting analysis as developed to detect cohesin DNA entrapment, we could clearly demonstrate the entrapment of a CEN plasmid by Smc5/6, similar to cohesin but without leading to cohesion of plasmid DNA (**Fig 6**). A control strain lacking one of the 6 cysteines for cross-linking of Smc5/6 did not show DNA entrapment.

**Figure 6:**
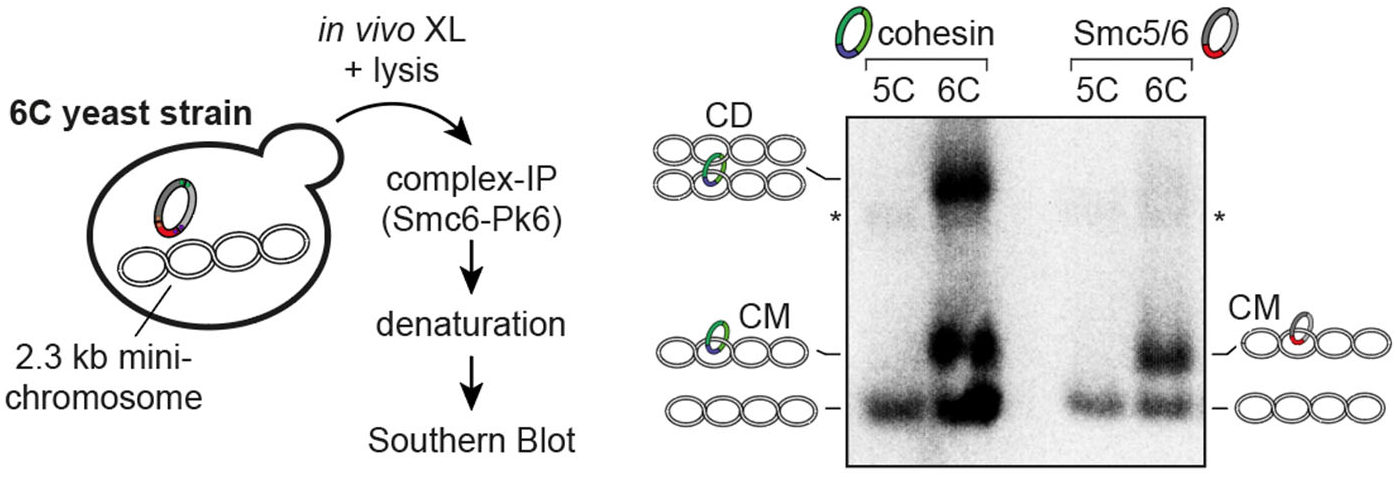
Smc5/6 entraps DNA *in vivo*. The scheme on the left shows an outline of the procedure. Results from Southern Blotting on the right show that cross-linkable versions (6C) of both cohesin and Smc5/6 entrap the minichromosome (catenated monomers; CM), but only cohesin does so in a cohesive manner (catenated dimers, CD). Control strains lacking one of the ring cysteines (5C) abolish entrapment in both cases.

## Discussion

### DNA entrapment by SMC complexes

Many of the diverse SMC functions in genome maintenance and chromosome organization are thought to arise directly from a DNA motor activity ^1^. Revealing the DNA topology underlying loop extrusion and translocation is a prerequisite for a basic understanding of SMC activity. We show here that purified preparations of Smc5/6 holocomplexes quickly and effectively entrap circular DNA substrates in a topological manner. This reaction is strictly ATP- and loader-dependent, and DNA entrapment is also detectable *in vivo*. We reveal the DNA entry gate and characterize the trajectory of DNA into the Smc5/6 ring showing that DNA loading involves a looped DNA substrate and eventually leads to the capture of a DNA segment by Smc5/6.

While the notion of DNA entrapment was challenged again recently by single-molecule imaging that apparently showed the bypass of very large obstacles on DNA by translocating cohesin and condensin ^5^, we argue based on our new findings and previous reports that DNA entrapment must not be discounted. Earlier studies measuring salt-stable DNA binding are in support of DNA entrapment by cohesin, condensin and Smc5/6, but they did go short of revealing the mode architecture of the hexameric Smc5/6 coreof DNA entrapment or the DNA-containing compartment ^16,20,56–58^. Experiments with entire bacterial chromosomes have provided strong evidence for DNA entrapment in the SK ring (and the K but not the S compartment) of Smc-ScpAB and MukBEF but did not reveal the mode of entrapment due to the complexity of the DNA substrate (including branches on the replicating chromosome) ^27,41^. Cohesin SK rings are known to support DNA entrapment *in vivo* at least in the context of sister chromatid cohesion ^26^. Single-molecule imaging experiments with cohesin however suggest that loop extrusion might not require cohesin ring opening, implying entrapment of a DNA loop or external DNA association rather than topological DNA entrapment ^2^. Entrapment of a DNA loop has recently also been proposed for the SK ring of yeast condensin based on co-isolation experiments after site-specific cross-linking ^50^.

Our data clearly demonstrate the topological DNA entrapment by Smc5/6 *in vivo* and elucidate its reliance on Nse5/6 *in vitro*. Nse5/6 has previously been implicated in chromosome loading *in vivo* by single molecule tracking in fission yeast ^32^. Curiously, Nse5/6 is dispensable for DNA loop extrusion by Smc5/6 *in vitro*, and actually hinders it ^34^. To reconcile these observations, one may invoke that Nse5/6 converts dynamic loop extruding Smc5/6 complexes into more stably bound DNA-entrapping complexes. The latter may (or may not) support DNA translocation ^54^. The functions of all these states and activities remain to be discerned.

### The loader opens the neck gate for DNA entry

Strict topological entrapment (as previously documented for cohesin and here for Smc5/6) requires the passage of one annular particle through an opening in another. Here, we unequivocally demonstrate that the neck gate serves as major and likely only entry gate in Smc5/6. This conclusion is based on an Smc6-Nse4 fusion protein, in which the linker does not block DNA loading but entraps the incoming DNA *in flagrante delicto* in an artificial linker compartment. The identity of the entry gate is furthermore supported by its efficient opening upon contact of the Smc5/6 hexamer with the loader Nse5/6, and by the lethality caused by the relevant fusion *in vivo* in budding yeast ^17^. Other possible gates do not seem to contribute to DNA entrapment in the reconstituted reaction [as judged by the absence of DNA entrapment by complexes harboring the Smc6-Nse4 fusion protein (as wells as cysteines for cross-linking the hinge and Smc5/Nse4)]. The situation appears more complicated in other SMC complexes. The neck gate is known to support DNA exit in cohesin ^60–64^ and has also been implicated in DNA entry in cohesin ^47,58^ as well as condensin ^65^. A recent paper demonstrated that cohesin DNA entry can be supported by the hinge and by the neck gate *in vitro*, however, with entry only via the hinge being dependent on the loader protein Scc2 ^53^, possibly suggesting that the hinge is the main or physiological gate for DNA entry, in contrast to what we observed here for Smc5/6. Moreover, sister chromatid cohesion can be built by cohesin complexes harboring an Smc3-Scc1 ‘neck gate’ fusion protein. This fusion protein also supports DNA entrapment by cohesin *in vivo* (when combined with cysteines for cross-linking of the other ring interfaces, unlike observed with Smc5/6 in **Fig 5**) ^26,59,66^ again implying fundamental differences with Smc5/6. A recent study in contrast to earlier reports suggested that condensin loading does not require an entry gate at least for DNA loop extrusion.

We find that the association with the loader destabilizes the contact between Smc6 and Nse4. Knowing that the loader intercalates between Smc5 and Smc6 heads ^19,20^ and that the Nse4 middle part is rather short, we propose that steric constraints prevent Nse4 from being attached to both SMC proteins simultaneously when the loader is also bound, resulting in the detachment of the least stable interface. DNA binding might reverse the effect of the loader, by evicting it from between the SMC heads and enabling productive head engagement (and ATP hydrolysis) ^20^ as well as ring re-closure by Smc6/Nse4 association. Assuming that the presence of DNA keeps the neck gate shut also during subsequent ATP hydrolysis cycles, this would result in a stable DNA association possibly facilitating DNA translocation over extended periods of time. The neck gate interface also appears labile in purified preparations of cohesin and condensin (sub-)complexes with the binding partners detaching from one another upon ATP-head engagement ^65,67^ [or incubation with cohesin unloading factors ^68^]. A similar reaction has been proposed for fission yeast Smc5/6 based on yeast-two-hybrid experiments ^69^. Neck gate opening upon ATP binding and head engagement, however, diametrically contrasts with our finding on budding yeast Smc5/6, where ATP-dependent head engagement leads to closure rather than opening of the gate (**Fig 7**). Whether these mechanistic differences have biological consequences will be important to work out.

**Figure 7:**
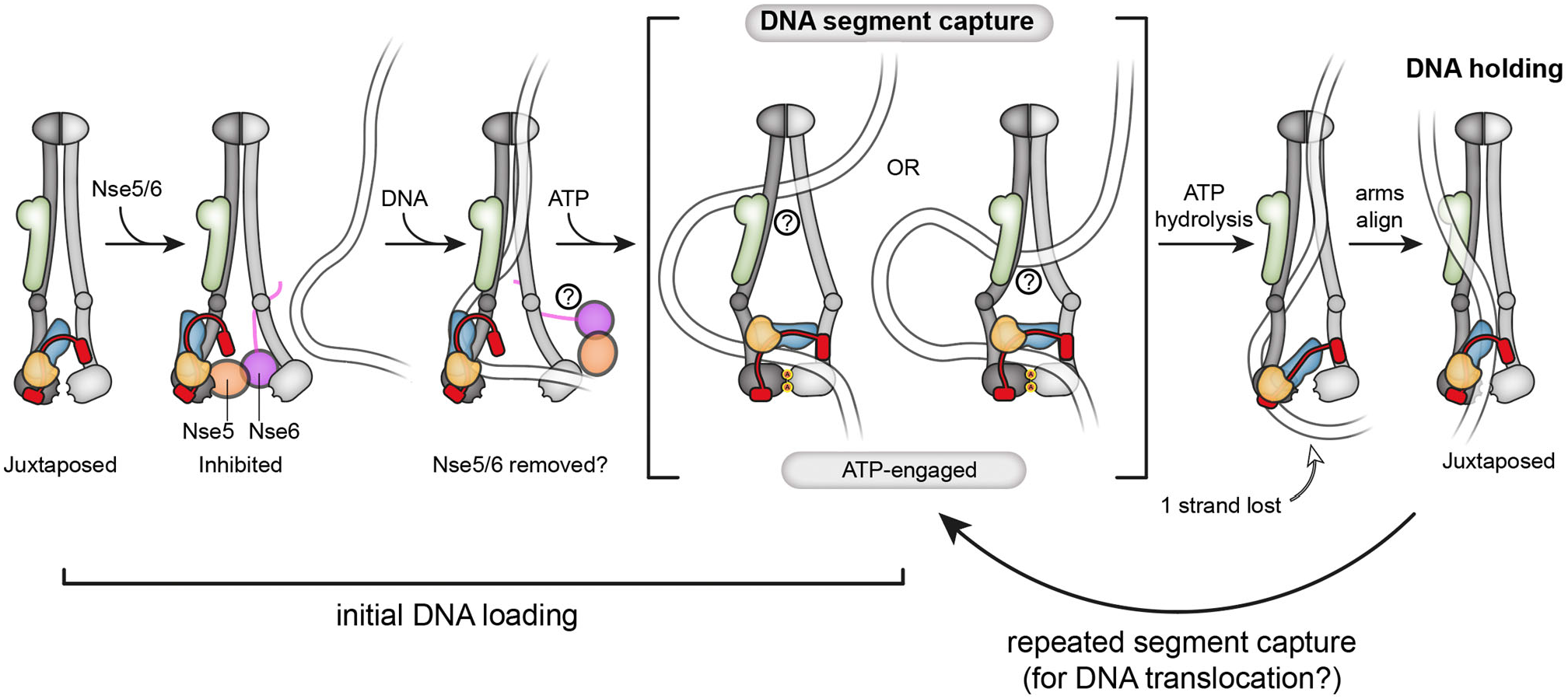
Model for chromosome loading and DNA translocation by alternating between a holding and a segment-capture state. Schematic representation of the main findings presented in this study. The segment-capture state (middle) is an essential intermediate in both DNA loading (left) and DNA translocation (right). Note that two possibilities for this state are indicated, differing in their degree of arm opening towards the hinge.

### DNA segment capture for loading as well as translocation

We propose that DNA loading involves DNA segment capture as an essential intermediate. We find that head/DNA binding is required for gate closure as well as for DNA entrapment. The head/DNA contacts conceivably hold a bent DNA segment in place during loading and further DNA contacts guide the kleisin subunit around DNA to ensure entrapment of a DNA double helix in the SK ring (**Fig 2C**). Of note, a model proposed for cohesin DNA loading via the neck gate resembles our notion of DNA segment capture ^47^. However, DNA loading via the cohesin neck gate has since been shown to work regardless of the absence or presence of the loading-clamping subunit Scc2, thus challenging this notion ^53^.

Why would DNA loading of SMC complexes involve such an intricate structure rather than rely on entrapping a single DNA double helix? DNA segment capture has first been proposed to explain DNA translocation and DNA loop extrusion by bacterial Smc-ScpAB and other SMC complexes ^36,54^ (**Fig 7**). We envisage that the segment-capture states during loading and during translocation/loop extrusion are structurally related. The notion of DNA segment capture mediating DNA loading as well as translocation explains how the very same ATPase core supports two distinct biochemical reactions and thus provides a unified framework for SMC ATPase functions, possibly also applicable to other SMC-related processes. The segment-capture state is consistent with observations on the bacterial SMC complexes Smc-ScpAB and MukBEF made by *in vivo* cross-linking and chromosome isolation as well as a cryo-EM structure of the MukBEF complex encompassing two separate DNA molecules ^38,41^. If true, the similarities in DNA topology are remarkable, especially when considering the large evolutionary distances between the three SMC complexes. Notably, the entrapment results were obtained with cysteines reporting distinct ATPase states (*i*.*e*. predominantly the juxtaposed and the ATP-engaged state for Smc-ScpAB and Smc5/6, respectively; in case of MukB the cysteines did not strongly discriminate between the states), indicating that DNA topology does not change after initial loading and that the entry gate may remain shut during subsequent ATP hydrolysis cycles (**Fig 7**). Recent data suggest that loop extrusion by condensin does not require the opening of a dedicated DNA entry gate and may therefore not involve the same loading intermediate ^50,70^. Subsequent steps of loop extrusion however could still proceed via DNA segment capture ^54^, which is consistent with the entrapment of DNA in the S^up^ compartment in condensin^50^.

Future experiments will have to elucidate how exactly DNA entrapment by Smc5/6 is linked to its function and its activities as translocator and loop extruder. While a segment capture-type mechanism explains our new findings, other suitable scenarios may exist or emerge in the future.

## Acknowledgements

We thank James Collier, Kim Nasmyth, and Madhusudhan Srinivasan for guidance and sharing reagents for the plasmid co-entrapment experiments. We are grateful to Serge Pelet and the Pelet lab for advice and for materials and reagents for genetic engineering in yeast and to Frank Bürmann, Markus Räschle, and members of the Gruber lab for comments on the manuscript and helpful discussions. This work was supported by the Swiss National Science Foundation (310030L_170242), the European Research Council (Horizon 2020 ERC CoG 724482) to S.G.

## Materials and Methods

### Cloning of expression plasmids

Plasmids containing multiple expression cassettes for subunits of the *S*. cerevisiae Smc5/6 complex were cloned as described in ^20^. Transcription of Smc5-containing cassettes was regulated with a tac-promoter and lambda-terminator, while the other subunits were transcribed by T7 promoters and terminators. A list of all expression plasmids used in this study can be found in Table S1.

### Protein expression and purification

All proteins and protein complexes described in this study were expressed in *E. coli* (DE3) Rosetta transformed with either a single plasmid or a combination of two plasmids. Table S2 lists plasmid details about all described complexes. For all purifications, 1 liter of the strain carrying the desired plasmid(s) was grown in TB-medium at 37°C to an OD(600nm) of 1.0 and the culture temperature was reduced to 22°C. Expression was then induced with IPTG at a final concentration of 0.4 mM and allowed to proceed overnight (typically for 16 hours). All Smc5/6 complexes and the Nse5/6 dimer were purified following a published procedure described in detail in ^20^.

### Cloning of yeast plasmids

Coding sequences for wildtype or mutant Smc6, Smc5, or Nse4 were cloned between their respective endogenous upstream and downstream regulatory sequences by Golden Gate cloning into centromeric acceptor plasmids pCEN(URA) or pCEN(TRP). N- or C-terminal tags were added as indicated. For plasmids containing two loci, we combined the individual cassettes by an additional Golden Gate Assembly step. A list of all yeast plasmids used in this study can be found in Table S3.

### Creation of yeast strains

All strains in this study were created in the W303 background. Functional assays to investigate mutants of Smc6, Smc5, and Nse4 were carried out using plasmid shuffling ^55^. To generate suitable strains, we first deleted the respective loci in a diploid strain then transformed the resulting strain with a pCEN(URA) plasmid containing the wild-type locus. Sporulation of this strain allowed us to isolate haploid offspring with the locus on the pCEN(URA) as the sole source for the respective protein. For double shuffling strains, both loci were deleted in the diploid and re-introduced on the pCEN(URA) plasmid. Haploid shuffling strains were transformed with pCEN(TRP) plasmids containing either wild-type or mutant version of the respective loci. A list of all yeast strains used in this study can be found in Table S4.

### Plasmid shuffling assays

Haploid shuffling strains containing the wild-type locus on pCEN(URA) and another wild-type or mutant locus on pCEN(TRP) were grown on plates lacking uracil and tryptophan. Single colonies were inoculated in minimal medium lacking only tryptophan and grown for 18-24 hours at 30°C. Four 10-fold dilutions were then prepared in water, and 2 µl of the 5 samples (undiluted culture and the 4 dilutions) were spotted on one plate lacking uracil and tryptophan, and on another containing all amino acids as well as 1 mg/ml 5-FOA (selecting for cells that had maintained or lost the pCEN(URA) plasmid, respectively). Plates were incubated at 30°C and pictures were taken at suitable time points (between 36 h and 60 h) to score growth of strains containing mutant versions of the respective Smc5/6 components.

### Site-specific BMOE crosslinking *in vitro*

Smc5/6 hexamers with or without indicated cysteines were diluted to a final concentration of 0.5 µM in ATPase buffer (10 mM HEPES-KOH pH 7.5, 150 mM KOAc, 2 mM MgCl_2_, 20 % glycerol) in a total volume of 30 µl. In reactions containing the Nse5/6 dimer, this complex was added in a 1.25 x molar excess (0.625 µM). For reactions containing ATP and/or plasmid DNA (25 kbp), these ligands were added at a final concentration of 2 mM and 5 nM, respectively. The circular or linear plasmid substrate was produced as described in ^20^. Protein and substrates were incubated for 5 min at RT after mixing, and BMOE was then added at a final concentration of 1 mM. After 45 seconds of incubation, dithiothreitol (DTT) was added at a final concentration of 10 mM to stop the reaction. For post-XL treatment with benzonase, 1 µl of the concentrated enzyme stock (750 U/µl) was added to the tube and incubated for 15 min at RT. Samples were mixed with SDS gel-loading dye, heated to 80°C for 15 min, and then analysed on Novex WedgeWell 4-12 % Tris-Glycine Gels or 3-8 % Tris-Acetate gels (Invitrogen). Tris-Glycine gels were run at 180 V for 1 hour at room temperature, Tris-Acetate gels were run at 4°C for 3 hours at 30 mA. Gels were fixed for 1-2 h in gel fixing solution (50 % ethanol, 10 % acetic acid) and stained overnight using Coomassie staining solution (50 % methanol, 10 % acetic acid, 1 mg/ml Coomassie Brilliant Blue R-250). Gels were destained in destaining solution (50 % methanol, 10 % acetic acid), and then rehydrated and stored in 5 % acetic acid. Quantification of bands in scanned gel images was done using Fiji ^71^.

### Analysis of salt-stable DNA association

These assays were carried out as described in (Taschner et al 2021) with modifications. 100 µl reactions in ATPase buffer (10 mM HEPESKOH pH 7.5, 150 mM KOAc, 2 mM MgCl_2_, 20 % glycerol) were set up containing combinations of the following components at the indicated concentrations: 600 nM of Smc5/6 hexameric complex (wildtype or mutant), 900 nM of Nse5/6 complex, 2 mM nucleotide, and 3 µg plasmid (pSG4418, 2.8 kbp). After incubation for 10 minutes at room temperature, 500 µl of ice-cold high-salt buffer (20 mM Tris pH 7.5, 1000 mM NaCl) were added and the mixture was incubated with 20 µl of StrepTactin Sepharose HP (GE Healthcare) for 45 minutes to pull out proteins via a C-terminal TwinStrep tag on Smc6. Beads were harvested by centrifugation (700g, 2 min) and washed twice with 1 ml of high salt buffer. The bound material was then eluted with a buffer containing 20 mM Tris pH 7.5, 250 mM NaCl, and 5 mM desthiobiotin. Aliquots of the eluate were either supplemented with 6 x gel loading dye containing SDS (Thermo Scientific), heated to 65° for 10 minutes, and analyzed by agarose gel electrophoresis (1 % in 0.5 x TBE) to check the DNA content, or with 2 x SDS-gel loading dye, heated to 95°C for 10 minutes, and analyzed by SDS-PAGE to visualize eluted proteins.

### Topological DNA binding assays

Analysis of topological DNA binding was performed following a protocol described in ^48^ with following modifications. For clear detection of ‘DNA laddering’ a small plasmid (1800 bp, pSG6085) was used at a final concentration of 15 nM and incubated with indicated concentrations of proteins in the presence or absence of 1 mM ATP. After incubation at RT (for 2 minutes except in the case of the time course shown in **Fig 2C**), BMOE was added to a final concentration of 1 mM. Crosslinking was allowed to proceed for 30 seconds at RT and then stopped by the addition of 10 mM DTT. For post-XL treatment with 3C protease, 1 µl of a stock of the home-made enzyme (concentration around 1 mg/ml) was added to the tube and incubated for 20 minutes at RT. The samples were supplemented with 6x loading dye with SDS (ThermoFisher) and heated for 20 minutes at 70°C. 10 µl aliquots were separated on 1 % agarose gels containing 0.03 % SDS and 1 µg/ml ethidium bromide. Gels were run at RT for 3 hours at 5 V/cm.

### Mini-chromosome entrapment assays

Entrapment of mini-chromosomes *in vivo* in budding yeast was performed folliwng a procedure previously used for cohesin (Srinivasan, Cell paper 2018?). Briefly, strains expressing a cross-linkable versions (‘6C’) of cohesin or Smc5/6 and containing a TRP mini-chromosome were grown together with control strains lacking one of the ring cysteines (‘5C’) overnight in medium lacking tryptophane. When the cultures reached an OD(600nm) of around 0.5, 40 ml (20 ODs) of the cultures were harvested by centrifugation (3000 rpm, 3 min), and the pellets were washed once with 25 ml ice-cold PBS and subsequently transferred to screw cap tubes. Pellets were then resuspended in 500 ml of ice-cold PBS. 30 µl of a 150 µM BMOE solution in DMSO were added and crosslinking was allowed to proceed for 6 minutes on ice. The cells were then pelleted, the supernatant was discarded, and cell pellets were snap-frozen in liquid nitrogen for storage at −80°C. For lysis, 700 µl of lysis buffer (50 mM HEPES-KOH pH 7.5, 100 mM KCl, 0.05 % Triton X-100, 0.025

% NP-40, 10 mM Na-citrate, 25 mM Na-sulfite) freshly supplemented with a cOmplete protease inhibitor tablet (Roche) and 1 mM PMSF were added to the pellet together with acid-washed glass beads (425-600 µm), and cells were opened using bead-beating with a FastPrep 24 system (MPBio; 3 x 1 min with 5 minute breaks on ice). The lysate was clarified by centrifugation at 14000 g for 10 min at 4°C, incubated with mouse monoclonal anti-V5 antibody (BioRad) for 90 min at 4°C to bind the C-terminal Pk6-epitope on Smc6, and complexes were then captured with DynaBeads Protein-G for another 90 min at 4°C. Beads were washed twice with 1 ml of wash buffer (10 mM Tris-HCl pH 8.0, 250 mM LiCl, 0.5 % NP-40, 0.5 % Na-deoxycholate, 1 mM EDTA) and once with 1 ml of TE, and bound material was then eluted with 1x DNA loading dye in TE supplemented with 1 % SDS and pre-heated to 65°C. The obtained material was separated on a 0.8 % agarose gel (in 1 x TAE) overnight at 4°C at 1.5 V/cm, and DNA was then transferred to Hybond-N+ membranes by Southern Blotting using alkaline transfer. Mini-chromosomes were detected with a probe against the TRP marker, which was labelled with alpha-32P-ATP using the Prime-It II Random Primer Labelling Kit (Agilent) according to the manufacturer’s instructions. Membranes were exposed to a Phosphor Storage Screen (Fujifilm) and signals were detected with a Typhoon Scanner (Cytiva).

### ATPase assays

Analysis of ATPase activity of selected Smc5/6 complexes were carried out exactly as described in ^20^.

**Figure S1:**
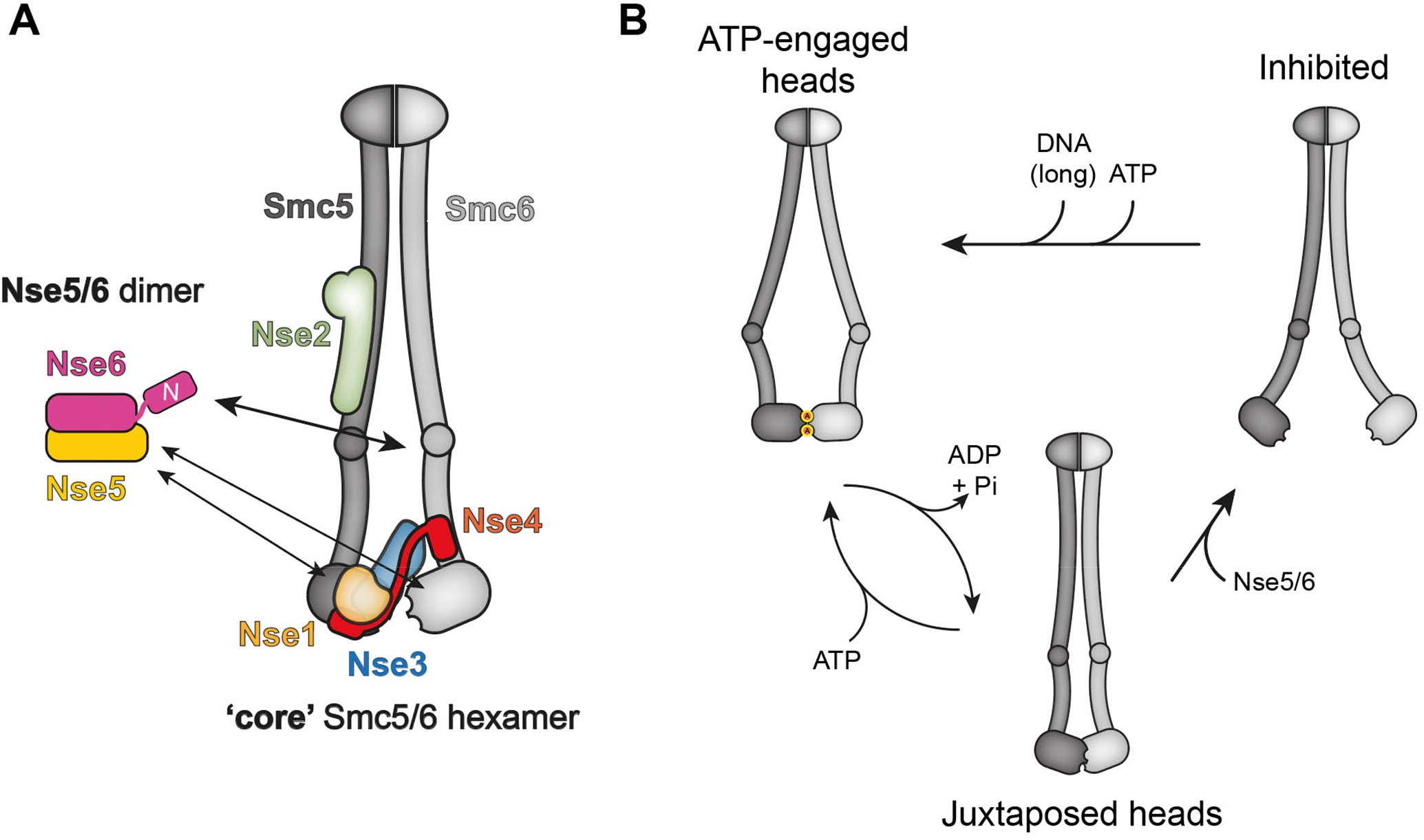
Schematics of Smc5/6 complex architecture. **(A)** Overall architecture of the hexameric Smc5/6 core complex and the Nse5/6 dimer (adapted from ^20^). For simplicity the Nse5/6 complex is shown separately with multiple arrows denoting various contact points with the core complex. (B)Simplified schematics focusing only on changes in Smc5/6 dimer architecture during the ATPase cycle and upon binding to the Nse5/6 loader and the DNA substrate.

**Figure S2:**
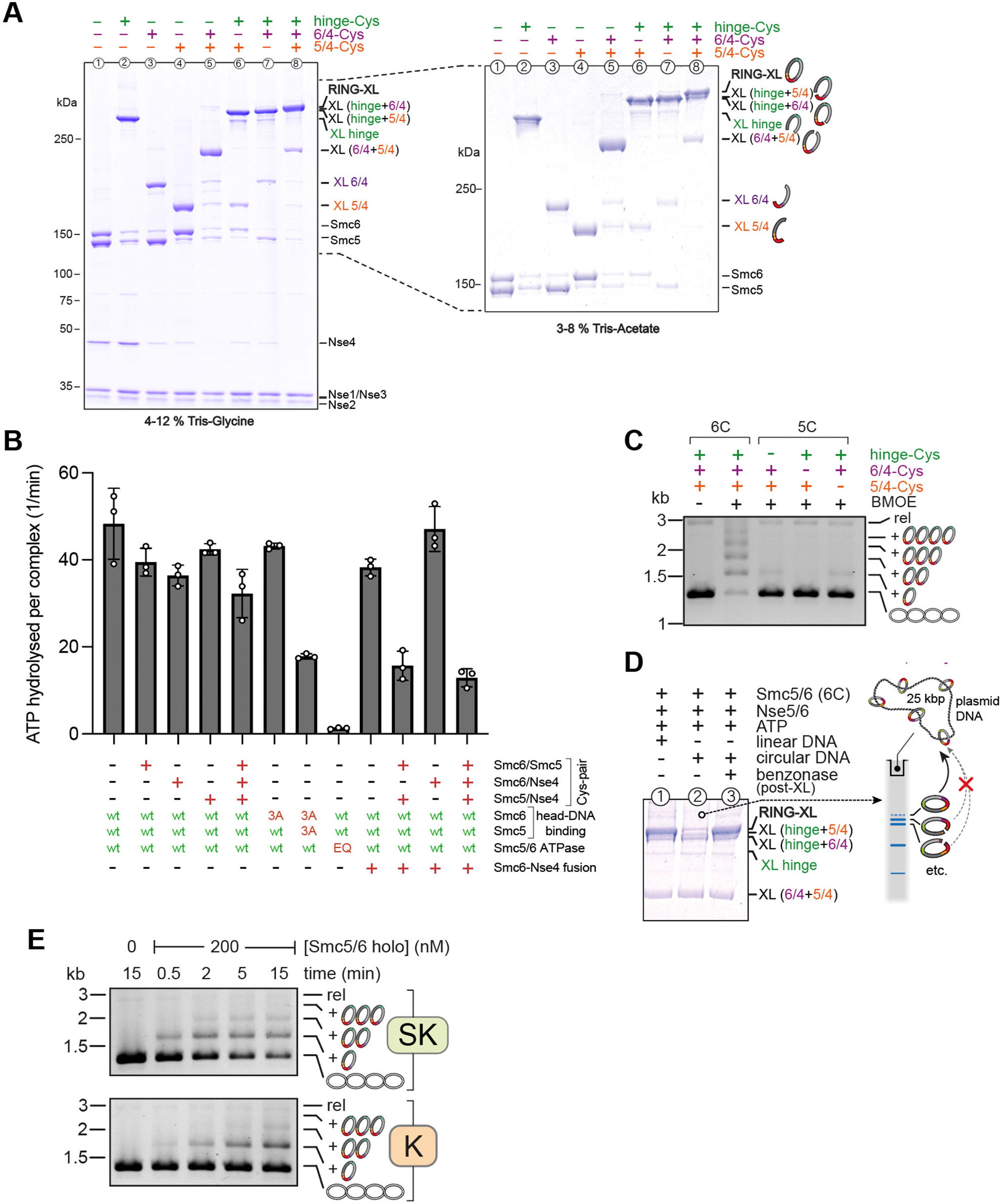
Analysis of Smc5/6 complexes harboring multiple engineered cysteines. **(A)** Cross-linking of Smc5/6 hexamers harboring multiple engineered cysteines as indicated. Identification of covalently closed ring species and intermediary cross-linking products by protein gel analysis on 2 types of gels (left: 4-12 % gradient gel, right: 3-8 % gradient gel) for proper separation of small and large proteins and detection by Coomassie staining. Note the presence of small amounts of ring species in two of the three control samples lacking one of the six cysteines (lanes 5 and 7), indicating minor off-target cross-linking. **(B)** ATPase activity assays with selected hexameric Smc5/6 complexes relevant for this study. Combinations of certain modifications lead to a reduction of ATPase activity. Error bars show standard deviations from technical triplicates (n=3). **(C)** Control experiments for entrapment experiments shown in Fig 1 with preparations lacking a selected cysteine (5C). Note that low levels of co-entrapment detected with two of the three 5C samples are likely explained by weak off-target cross-linking. **(D)** Under conditions promoting DNA entrapment the ring species is visible in a protein gel in the presence of a linear (lane 1) but not a circular (lane 2) DNA substrate, presumably due to co-retention of the ring species with circular DNA in the loading well (see scheme on the right). The ring species re-appears after digestion of the circular substrate (lane 3). (E) Time-dependence of DNA entrapment in the SK ring (top panel) and the K compartment (bottom panel). Cross-linker was added to a sample aliquot at the indicated time points.

**Figure S3:**
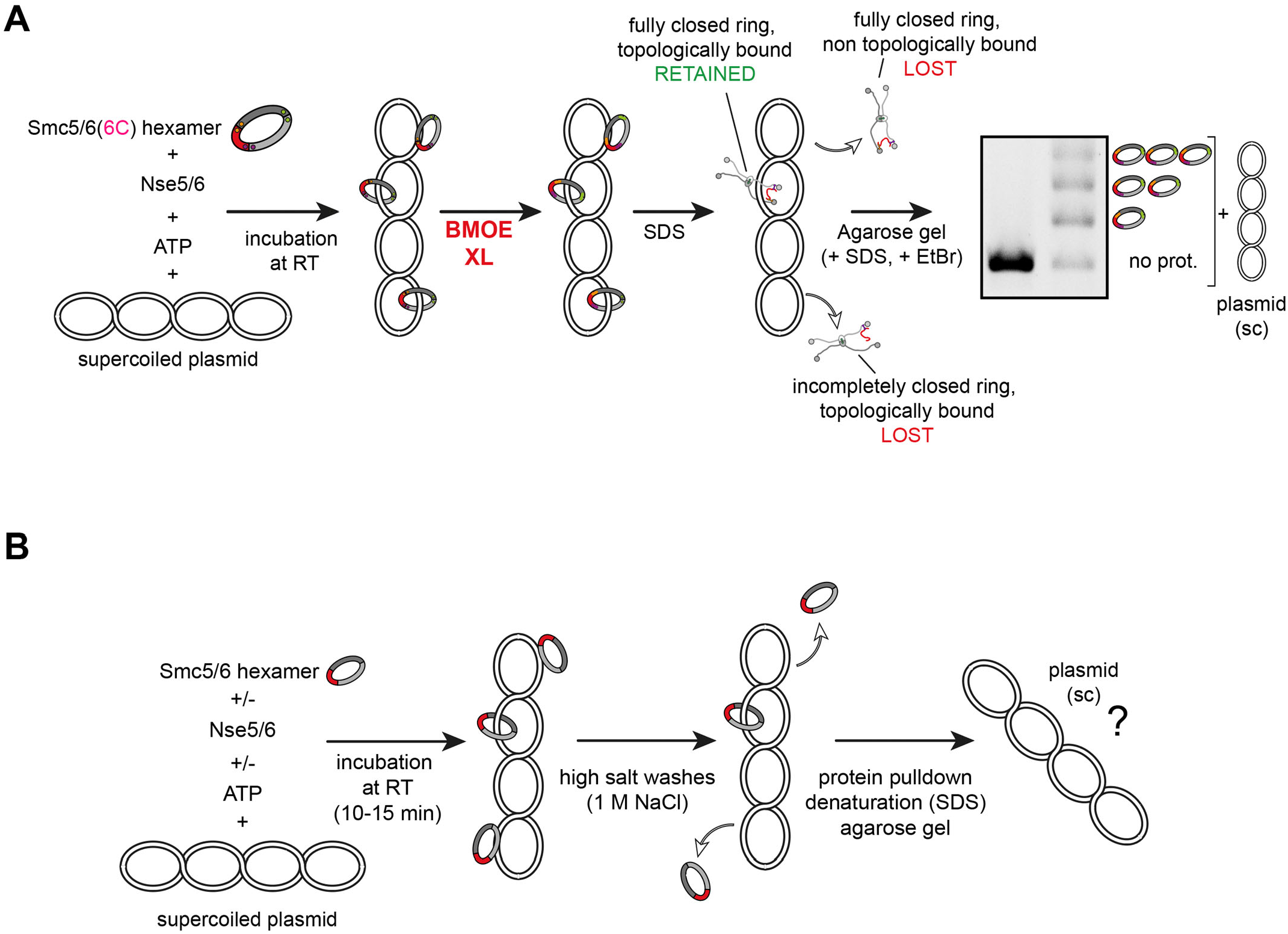
Schematic overview of assays used to examine the nature of Smc5/6 association with its DNA substrate. **(A)** In the topological loading assay, a cross-linkable version of the Smc5/6 hexamer (‘6C’) is incubated with a small supercoiled (‘sc’) plasmid substrate in the absence or presence of ATP and Nse5/6. Upon cross-linking with BMOE and protein denaturation (‘SDS’) only fully cross-linked rings that were topologically associated with the substrate are retained, leading to a characteristic laddering pattern in agarose gels. **(B)** The salt-stable binding assay does not involve cross-linking. Complexes are first incubated with the substrate under low-salt conditions before the salt concentration is increased to 1 M NaCl by buffer changes. Smc5/6 complexes are immobilized on beads using a Twin-Strep tag on Smc6, and co-purified plasmid substrate is examined on an agarose gel.

**Figure S4:**
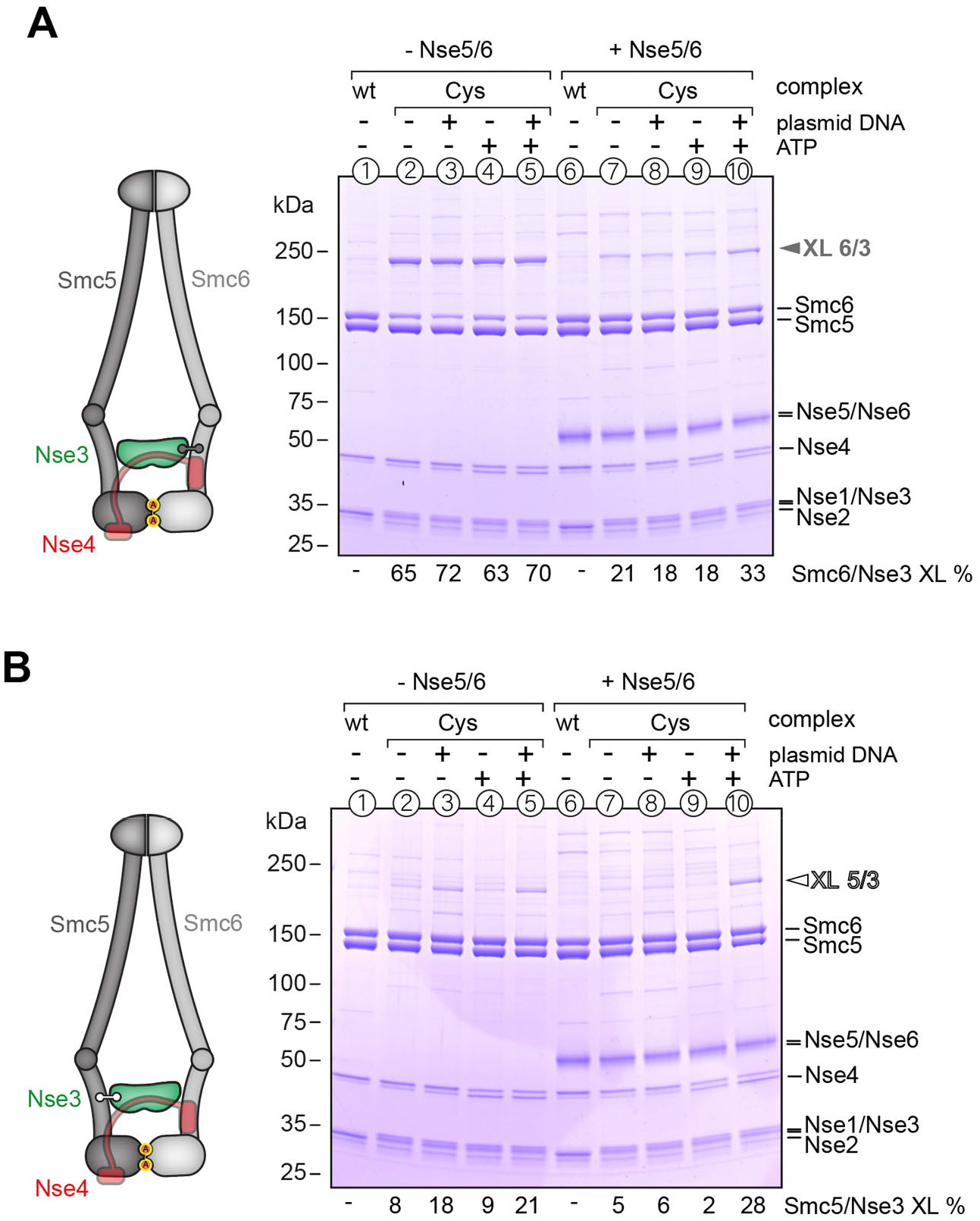
Analysis of Nse3 crosslinking to SMC arms. Cross-linking of purified Smc5/6 hexamers with cysteines at the Smc6/Nse3 interface **(A)** or at the Smc5/Nse3 interface **(B)** in the presence and absence of ligands. Detection of cross-linked species by SDS-Page and Coomassie staining. Numbers below the gel indicate the efficiency of the respective cross-link.

**Figure S5:**
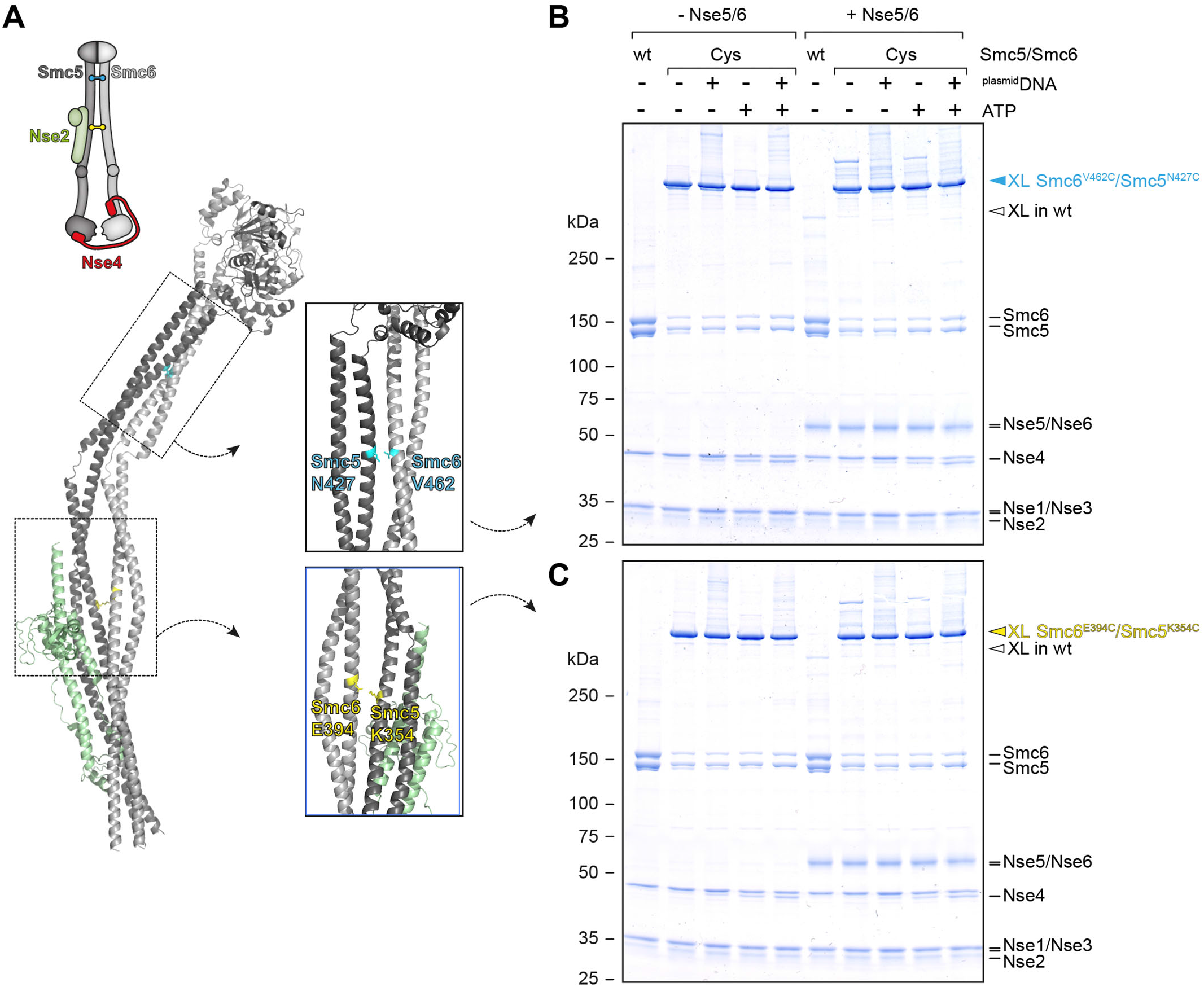
Head-distal Smc5/6 coiled-coil arms do not detectably open upon addition of Nse5/6, ATP, and/or DNA. **(A)** Model of an Smc5/Smc6/Nse2 complex obtained with AlphaFold-Multimer. The positions of two engineered cysteine pairs for inter-arm cross-linking are denoted. **(B, C)** Results from cross-linking experiments showing efficient arm alignment in all tested conditions. As in Fig 3 and fig S6.

**Figure S6:**
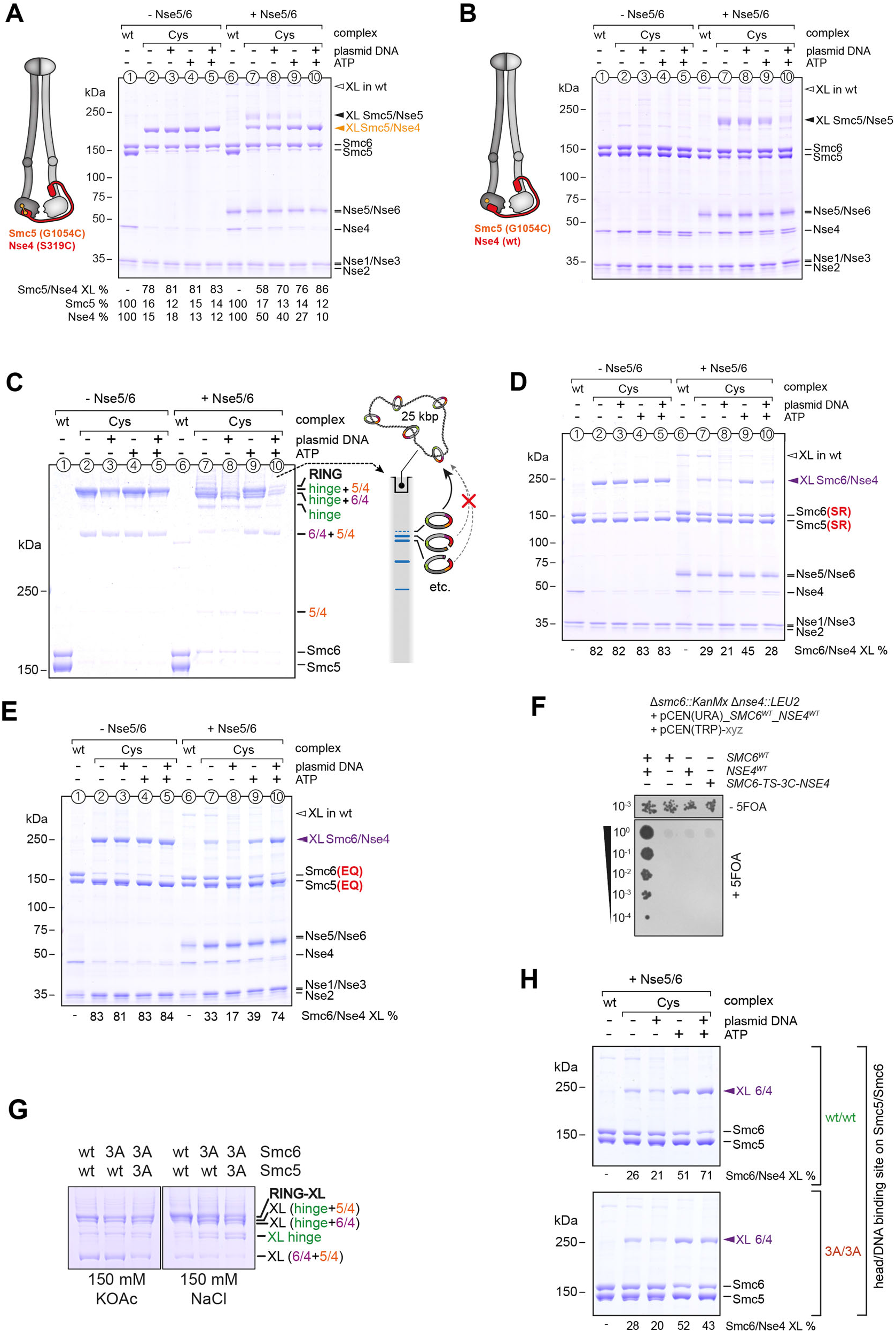
Cross-linking of selected SMC/kleisin interfaces. Similar to Fig 3 **(A)** The Smc5/Nse4 interface is efficiently cross-linked and does not detectably respond to the presence of ligands apart from weak off-target cross-linking between Smc5 and the loader subunit Nse5 [see (B)]. **(B)** Control reaction for (A) with protein samples lacking the engineered cysteine in Nse4 confirming off-target cross-linking of Smc5(G1054C) to Nse5. (B)Addition of the loader reduces abundance of the ring species due to gate opening. The pattern in the presence of ligands (ATP and plasmid DNA) mirrors the one obtained with the Smc6/Nse4 interface (Fig 3), except when loader, ATP and plasmid are added, presumably due to co-retention of ring species with circular DNA in the loading well (see scheme on the right). **(D)** As in Fig 3 but with Smc5 and Smc6 subunits carrying the signature motif head-engagement mutation (‘SR’). **(E)** As in Fig 3 but with Smc5 and Smc6 subunits carrying the Walker B ATP hydrolysis mutation (‘EQ’). **(F)** Fusion of Smc6 to Nse4 with the linker used in our *in vitro* assays is lethal in yeast. Plasmid shuffling assay as in **fig S7F**, but with a deletion mutant for both *smc6* and *nse4*. Adding back both wildtype genes separately, but not either of them alone or a fused version, restores viability. **(G)** Effect of mutating the head DNA binding interfaces on Smc5 and/or Smc6 on formation of a closed SMC/Kleisin (SK) ring. **(H)** Effect of mutating the head DNA binding interfaces on Smc5 and Smc6 on Smc6/Nse4 gate closure in the presence of the loader and various ligands.

**Figure S7:**
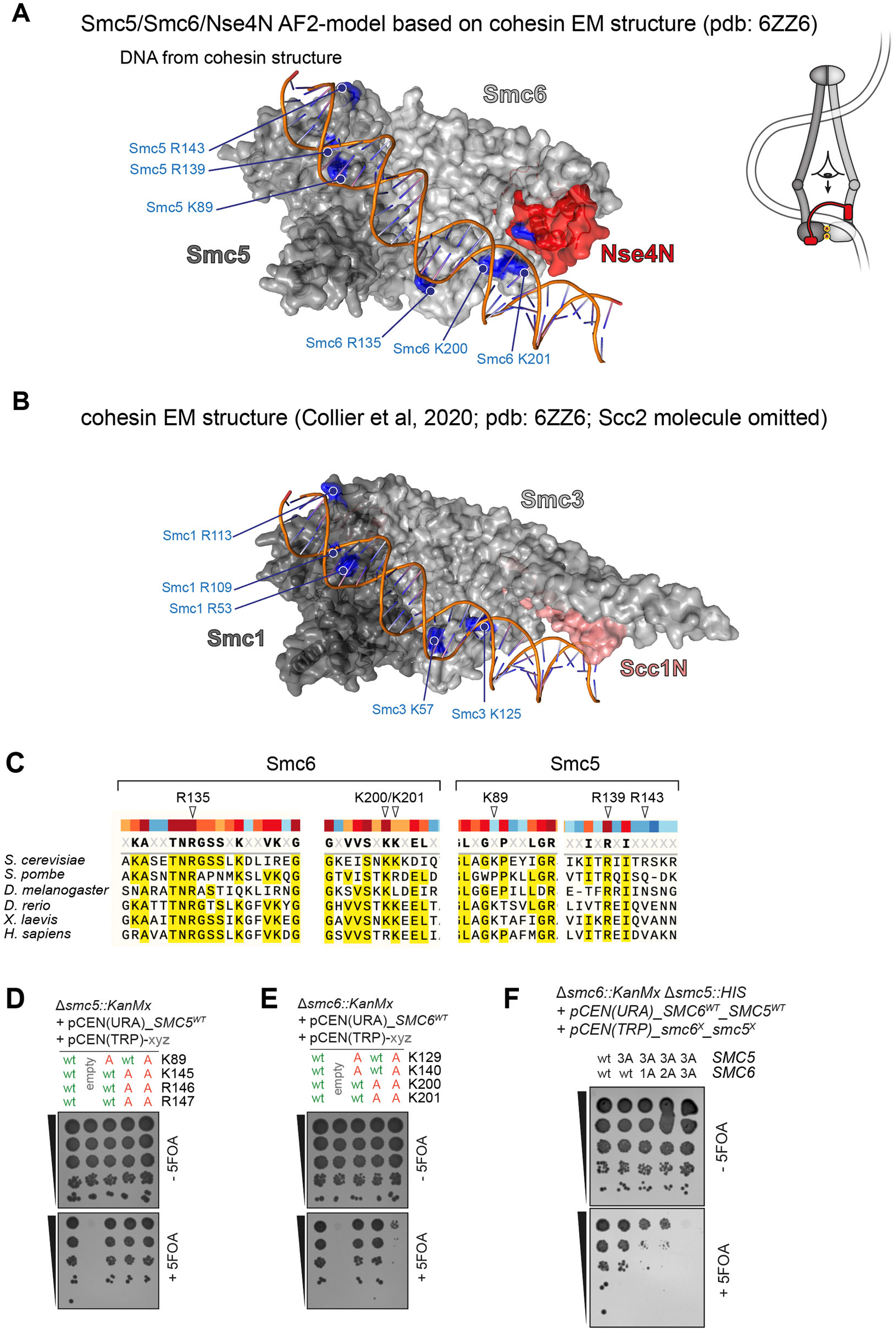
Identification of putative DNA binding residues on Smc6 and Smc5. **(A)** Model of ATP-engaged Smc6/Smc5 heads with the Nse4 N-terminus bound to the Smc6 neck. AlphaFold-Multimer models of Smc6/Nse4 and Smc5 were superimposed on their counterparts of the cohesin complex (pdb 6ZZ6, see panel B). The DNA molecule from the cohesin cryo-EM structure contacts several putative DNA binding residues on Smc6 and Smc5. **(B)** Similar view on top of engaged heads in the cohesin cryo-EM structure (pdb 6ZZ6) with DNA-interacting residues on Smc3 and Smc1 indicated. Note that the Scc2 molecule is not shown for simplicity. **(C)** Sequence alignment showing strong evolutionary conservation of examined residues in Smc6. Smc5 residues show weaker conservation consistent with results of functional assays shown in Fig 4. **(D)** Positively charged residues on the Smc5 head were mutated to alanine (‘A’) and the mutant alleles were tested for function by plasmid shuffling. As in Fig 4B but with residues selected based on a recent cryo-EM structure of Smc5/6 (Yu *et al*, 2022). **(E)** As in (D) but for residues on the Smc6 head. **(F)** Positively charged residues on Smc5 and Smc6 heads were mutated to alanines in combinations. As in Fig 4B and C.

**Figure S8:**
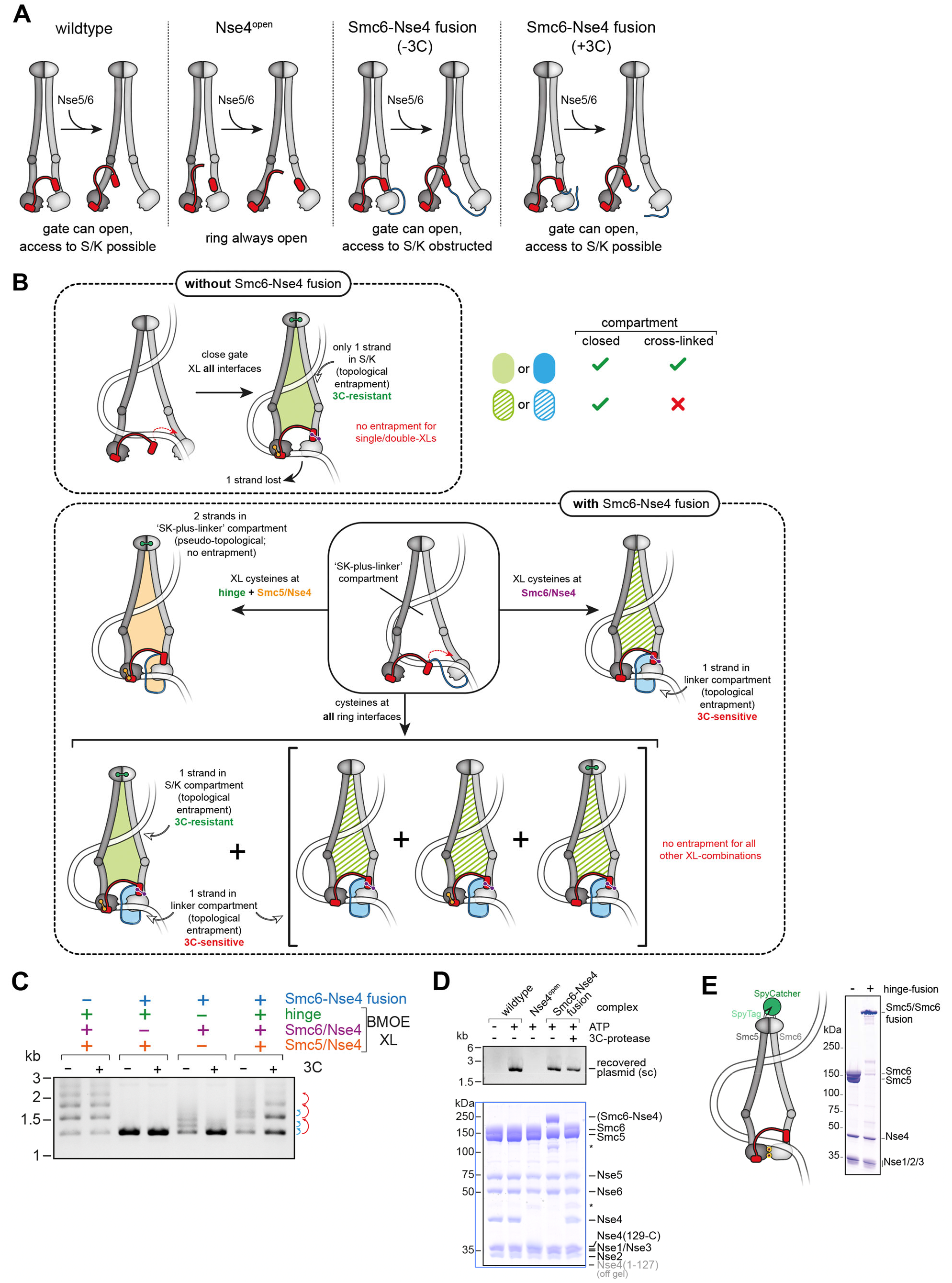
Entrapment assays involving Smc6-Nse4 fusion and BMOE cross-linking of interfaces. **(A)** Schematic representation of complexes used for experiments in Fig 5 and fig S8D. The ‘Nse4^open^’ complex has a split kleisin and thus has its SK ring permanently opened. The ‘Smc6-Nse4 fusion’ complex contains a peptide linker fusing the C-terminus of Smc6 with the N-terminus of Nse4. The linker contains a recognition site for the HRV 3C protease and can thus be opened by cleavage. **(B)** Schematic overview of entrapment in the absence and presence of the Smc6-Nse4 fusion. Without the fusion (top) during initial DNA segment capture the Smc6/Nse4 gate is opened by Nse5/6 (Smc5 and Smc6 arm distance is exaggerated for clarity). Upon gate closure and ATP hydrolysis, one of the loop strands is lost after escape between the disengaged heads, and the other strand becomes topologically entrapped in the SK ring. The green-filled area indicates the lumen of the ring compartment (SK) which is maintained even after protein denaturation due to cross-linking. (bottom) Scenarios for DNA entrapment by complexes with Smc6-Nse4 fusion (top row, middle panel) in combination with different cysteine pairs for cross-linking (other panels). Top row, left panel: Upon cross-linking of Smc5/Smc6 and Smc5/Nse4 interfaces an SK-plus-linker joint compartment becomes denaturation-resistant (orange-filled area) leading to DNA loop entrapment rather than DNA entrapment (detected in this assay). Top row, right panel: Cross-linking of only the Smc6/Nse4 interface leads to a denaturation-resistant linker compartment (blue-filled area) that topologically entraps one of the two DNA strands. The SK compartment is also formed and entraps the other DNA strand, but it is sensitive to denaturation (green-dashed area). Bottom row: Cross-linking of all three interfaces entraps one DNA strand in a denaturation-resistant SK compartment (green-filled area) and another in the linker compartment (blue-filled area), only the latter of which can be released by incubation with 3C protease. Incompletely cross-linked rings lacking the Smc5/Smc6 or Smc5/Nse4 cross-links (or both) entrap only the DNA strand in the linker compartment (in square bracket). **(C)** As in Fig 5C but with cleavage of the linker prior to mixing of samples for DNA entrapment. **(D)** Salt-stable DNA binding of Smc5/6 complexes with a split Nse4 protein (‘Nse4 open’) or linked Smc6 and Nse4 proteins (‘Smc6-Nse4’ fusion protein). As in Fig 3D. Pre-treatment with 3C protease cleaves the fusion linker peptide but does not alter DNA binding. Asterisks denote unspecific degradation products. **(E)** Insertion of a Spy-tag and Spy-Catcher into the Smc5 and Smc6 hinge-domains (see scheme on the left), respectively, leads to permanent, covalent fusion of the hinge domains as shown in the protein gel on the right.

## References

1. Yatskevich, S., Rhodes, J. & Nasmyth, K. Organization of Chromosomal DNA by SMC Complexes. Annu Rev Genet 53, 445–482 (2019).

2. Davidson, I.F. et al. DNA loop extrusion by human cohesin. Science 366, 1338–1345 (2019).

3. Ganji, M. et al. Real-time imaging of DNA loop extrusion by condensin. Science 360, 102–105 (2018).

4. Kim, Y., Shi, Z., Zhang, H., Finkelstein, I.J. & Yu, H. Human cohesin compacts DNA by loop extrusion. Science 366, 1345–1349 (2019).

5. Pradhan, B. et al. SMC complexes can traverse physical roadblocks bigger than their ring size. bioRxiv (2021).

6. Datta, S., Lecomte, L. & Haering, C.H. Structural insights into DNA loop extrusion by SMC protein complexes. Curr Opin Struct Biol 65, 102–109 (2020).

7. Higashi, T.L. & Uhlmann, F. SMC complexes: Lifting the lid on loop extrusion. Curr Opin Cell Biol 74, 13–22 (2022).

8. Aragon, L. The Smc5/6 Complex: New and Old Functions of the Enigmatic Long-Distance Relative. Annu Rev Genet 52, 89–107 (2018).

9. McDonald, W.H., Pavlova, Y., Yates, J.R., 3rd & Boddy, M.N. Novel essential DNA repair proteins Nse1 and Nse2 are subunits of the fission yeast Smc5-Smc6 complex. J Biol Chem 278, 45460–7 (2003).

10. Pebernard, S., McDonald, W.H., Pavlova, Y., Yates, J.R., 3rd & Boddy, M.N. Nse1, Nse2, and a novel subunit of the Smc5-Smc6 complex, Nse3, play a crucial role in meiosis. Mol Biol Cell 15, 4866–76 (2004).

11. Copsey, A. et al. Smc5/6 coordinates formation and resolution of joint molecules with chromosome morphology to ensure meiotic divisions. PLoS Genet 9, e1004071 (2013).

12. Xaver, M., Huang, L., Chen, D. & Klein, F. Smc5/6-Mms21 prevents and eliminates inappropriate recombination intermediates in meiosis. PLoS Genet 9, e1004067 (2013).

13. Torres-Rosell, J. et al. SMC5 and SMC6 genes are required for the segregation of repetitive chromosome regions. Nat Cell Biol 7, 412–9 (2005).

14. Torres-Rosell, J. et al. The Smc5-Smc6 complex and SUMO modification of Rad52 regulates recombinational repair at the ribosomal gene locus. Nat Cell Biol 9, 923–31 (2007).

15. Kegel, A. et al. Chromosome length influences replication-induced topological stress. Nature 471, 392–6 (2011).

16. Gutierrez-Escribano, P. et al. Purified Smc5/6 Complex Exhibits DNA Substrate Recognition and Compaction. Mol Cell 80, 1039–1054 e6 (2020).

17. Serrano, D. et al. The Smc5/6 Core Complex Is a Structure-Specific DNA Binding and Compacting Machine. Mol Cell 80, 1025–1038 e5 (2020).

18. Yu, Y. et al. Integrative analysis reveals unique structural and functional features of the Smc5/6 complex. Proc Natl Acad Sci U S A 118(2021).

19. Hallett, S.T. et al. Nse5/6 is a negative regulator of the ATPase activity of the Smc5/6 complex. Nucleic Acids Res 49, 4534–4549 (2021).

20. Taschner, M. et al. Nse5/6 inhibits the Smc5/6 ATPase and modulates DNA substrate binding. EMBO J 40, e107807 (2021).

21. Hallett, S.T. et al. Cryo-EM structure of the Smc5/6 holo-complex. Nucleic Acids Res (2022).

22. Lammens, A., Schele, A. & Hopfner, K.P. Structural biochemistry of ATP-driven dimerization and DNA-stimulated activation of SMC ATPases. Curr Biol 14, 1778–82 (2004).

23. Hopfner, K.P. Invited review: Architectures and mechanisms of ATP binding cassette proteins. Biopolymers 105, 492–504 (2016).

24. Hirano, M., Anderson, D.E., Erickson, H.P. & Hirano, T. Bimodal activation of SMC ATPase by intra- and inter-molecular interactions. EMBO J 20, 3238–50 (2001).

25. Burmann, F. et al. An asymmetric SMC-kleisin bridge in prokaryotic condensin. Nat Struct Mol Biol 20, 371–9 (2013).

26. Gligoris, T.G. et al. Closing the cohesin ring: structure and function of its Smc3-kleisin interface. Science 346, 963–7 (2014).

27. Wilhelm, L. et al. SMC condensin entraps chromosomal DNA by an ATP hydrolysis dependent loading mechanism in Bacillus subtilis. Elife 4(2015).

28. Palecek, J., Vidot, S., Feng, M., Doherty, A.J. & Lehmann, A.R. The Smc5-Smc6 DNA repair complex. bridging of the Smc5-Smc6 heads by the KLEISIN, Nse4, and non-Kleisin subunits. J Biol Chem 281, 36952–9 (2006).

29. Haering, C.H. et al. Structure and stability of cohesin’s Smc1-kleisin interaction. Mol Cell 15, 951–64 (2004).

30. Palecek, J.J. & Gruber, S. Kite Proteins: a Superfamily of SMC/Kleisin Partners Conserved Across Bacteria, Archaea, and Eukaryotes. Structure 23, 2183–2190 (2015).

31. Duan, X. et al. Structural and functional insights into the roles of the Mms21 subunit of the Smc5/6 complex. Mol Cell 35, 657–68 (2009).

32. Etheridge, T.J. et al. Live-cell single-molecule tracking highlights requirements for stable Smc5/6 chromatin association in vivo. Elife 10(2021).

33. Pebernard, S., Wohlschlegel, J., McDonald, W.H., Yates, J.R., 3rd & Boddy, M.N. The Nse5-Nse6 dimer mediates DNA repair roles of the Smc5-Smc6 complex. Mol Cell Biol 26, 1617–30 (2006).

34. Pradhan, B. et al. The Smc5/6 complex is a DNA loop extruding motor. bioRxiv (2022).

35. Oravcová, M. et al. The Nse5/6-like SIMC1-SLF2 Complex Localizes SMC5/6 to Viral Replication Centers. bioRxiv (2022).

36. Diebold-Durand, M.L. et al. Structure of Full-Length SMC and Rearrangements Required for Chromosome Organization. Mol Cell 67, 334–347 e5 (2017).

37. Soh, Y.M. et al. Molecular basis for SMC rod formation and its dissolution upon DNA binding. Mol Cell 57, 290–303 (2015).

38. Vazquez Nunez, R., Ruiz Avila, L.B. & Gruber, S. Transient DNA Occupancy of the SMC Interarm Space in Prokaryotic Condensin. Mol Cell 75, 209–223 e6 (2019).

39. Chapard, C., Jones, R., van Oepen, T., Scheinost, J.C. & Nasmyth, K. Sister DNA Entrapment between Juxtaposed Smc Heads and Kleisin of the Cohesin Complex. Mol Cell 75, 224–237 e5 (2019).

40. Lee, B.G. et al. Cryo-EM structures of holo condensin reveal a subunit flip-flop mechanism. Nat Struct Mol Biol 27, 743–751 (2020).

41. Bürmann, F., Funke, L.F.H., Chin, J.W. & Löwe, J. Cryo-EM structure of MukBEF reveals DNA loop entrapment at chromosomal unloading sites. Molecular Cell (2021).

42. Vazquez Nunez, R., Polyhach, Y., Soh, Y.M., Jeschke, G. & Gruber, S. Gradual opening of Smc arms in prokaryotic condensin. Cell Rep 35, 109051 (2021).

43. Yu, Y. et al. Cryo-EM structure of DNA-bound Smc5/6 reveals DNA clamping enabled by multi-subunit conformational changes. Proc Natl Acad Sci U S A 119, e2202799119 (2022).

44. Seifert, F.U., Lammens, K., Stoehr, G., Kessler, B. & Hopfner, K.P. Structural mechanism of ATP-dependent DNA binding and DNA end bridging by eukaryotic Rad50. EMBO J 35, 759–72 (2016).

45. Shi, Z., Gao, H., Bai, X.C. & Yu, H. Cryo-EM structure of the human cohesin-NIPBL-DNA complex. Science 368, 1454–1459 (2020).

46. Liu, Y. et al. ATP-dependent DNA binding, unwinding, and resection by the Mre11/Rad50 complex. EMBO J 35, 743–58 (2016).

47. Higashi, T.L. et al. A Structure-Based Mechanism for DNA Entry into the Cohesin Ring. Mol Cell 79, 917–933 e9 (2020).

48. Collier, J.E. et al. Transport of DNA within cohesin involves clamping on top of engaged heads by Scc2 and entrapment within the ring by Scc3. Elife 9(2020).

49. Lee, B.-G., Rhodes, J. & Löwe, J. Clamping of DNA shuts the condensin neck gate. bioRxiv (2021).

50. Shaltiel, I.A. et al. A hold-and-feed mechanism drives directional DNA loop extrusion by condensin. Science 376, 1087–1094 (2022).

51. Jumper, J. et al. Highly accurate protein structure prediction with AlphaFold. Nature 596, 583–589 (2021).

52. Evans, R. et al. Protein complex prediction with AlphaFold-Multimer. bioRxiv (2021).

53. Collier, J.E. & Nasmyth, K.A. DNA passes through cohesin’s hinge as well as its Smc3-kleisin interface. Elife 11(2022).

54. Marko, J.F., De Los Rios, P., Barducci, A. & Gruber, S. DNA-segment-capture model for loop extrusion by structural maintenance of chromosome (SMC) protein complexes. Nucleic Acids Res 47, 6956–6972 (2019).

55. Boeke, J.D., Trueheart, J., Natsoulis, G. & Fink, G.R. 5-Fluoroorotic acid as a selective agent in yeast molecular genetics. Methods Enzymol 154, 164–75 (1987).

56. Kanno, T., Berta, D.G. & Sjogren, C. The Smc5/6 Complex Is an ATP-Dependent Intermolecular DNA Linker. Cell Rep 12, 1471–82 (2015).

57. Cuylen, S., Metz, J. & Haering, C.H. Condensin structures chromosomal DNA through topological links. Nat Struct Mol Biol 18, 894–901 (2011).

58. Murayama, Y. & Uhlmann, F. Biochemical reconstitution of topological DNA binding by the cohesin ring. Nature 505, 367–71 (2014).

59. Gruber, S. et al. Evidence that loading of cohesin onto chromosomes involves opening of its SMC hinge. Cell 127, 523–37 (2006).

60. Chan, K.L. et al. Cohesin’s DNA exit gate is distinct from its entrance gate and is regulated by acetylation. Cell 150, 961–74 (2012).

61. Elbatsh, A.M.O. et al. Cohesin Releases DNA through Asymmetric ATPase-Driven Ring Opening. Mol Cell 61, 575–588 (2016).

62. Ouyang, Z. & Yu, H. Releasing the cohesin ring: A rigid scaffold model for opening the DNA exit gate by Pds5 and Wapl. Bioessays 39(2017).

63. Eichinger, C.S., Kurze, A., Oliveira, R.A. & Nasmyth, K. Disengaging the Smc3/kleisin interface releases cohesin from Drosophila chromosomes during interphase and mitosis. EMBO J 32, 656–65 (2013).

64. Beckouet, F. et al. Releasing Activity Disengages Cohesin’s Smc3/Scc1 Interface in a Process Blocked by Acetylation. Mol Cell 61, 563–574 (2016).

65. Hassler, M. et al. Structural Basis of an Asymmetric Condensin ATPase Cycle. Mol Cell 74, 1175–1188 e9 (2019).

66. Srinivasan, M. et al. The Cohesin Ring Uses Its Hinge to Organize DNA Using Non-topological as well as Topological Mechanisms. Cell 173, 1508–1519 e18 (2018).

67. Muir, K.W., Li, Y., Weis, F. & Panne, D. The structure of the cohesin ATPase elucidates the mechanism of SMC-kleisin ring opening. Nat Struct Mol Biol 27, 233–239 (2020).

68. Murayama, Y. & Uhlmann, F. DNA Entry into and Exit out of the Cohesin Ring by an Interlocking Gate Mechanism. Cell 163, 1628–1640 (2015).

69. Vondrova, L. et al. A role of the Nse4 kleisin and Nse1/Nse3 KITE subunits in the ATPase cycle of SMC5/6. Sci Rep 10, 9694 (2020).

70. Shaltiel, I.A. et al. A hold-and-feed mechanism drives directional DNA loop extrusion by condensin. bioRxiv, 2021.10.29.466147 (2021).

71. Schindelin, J. et al. Fiji: an open-source platform for biological-im-age analysis. Nat Methods 9, 676–82 (2012).

